# Heritability of the Mouse Brain Connectome

**DOI:** 10.1101/701755

**Authors:** Nian Wang, Robert J Anderson, David G Ashbrook, Vivek Gopalakrishnan, Youngser Park, Carey E Priebe, Yi Qi, Joshua T Vogelstein, Robert W Williams, G Allan Johnson

## Abstract

Genome-wide association studies have demonstrated significant links between human brain structure and common DNA variants. Similar studies with rodents have been challenging because of smaller brain volumes. Using high field MRI (9.4T) and compressed sensing, we have achieved microscopic resolution and sufficiently high throughput for rodent population studies. We generated whole brain structural MRI and diffusion connectomes for four diverse isogenic lines of mice (C57BL/6J, DBA/2J, CAST/EiJ, and BTBR) at spatial resolution 20,000 times higher than human connectomes. We derived volumes, scalar diffusion metrics, and estimates of residual technical error for 166 regions in each hemisphere and connectivity between the regions. Volumes of discrete brain regions had the highest mean heritability (0.71 ± 0.23 SD, *n* = 332), followed by fractional anisotropy (0.54 ± 0.26), radial diffusivity (0.34 ± 0.022), and axial diffusivity (0.28 ± 0.19). Connection profiles were statistically different in 280 of 322 nodes across all four strains. Nearly 150 of the connection profiles were statistically different between the C57BL/6J, DBA/2J, and CAST/EiJ lines.

## INTRODUCTION

Structural and functional magnetic resonance imaging (MRI) have enabled new ways to systematically study the influence of DNA sequence differences on brain architecture and behavior. Investigators have used both conventional family linkage studies and genome-wide association studies [1–5]. New DTI methods provide enhanced structural detail, scalar measures of tissue cytoarchitecture, and structural connections across the entire brain [5, 6].

Experimental genetic studies that have used many of these MRI methods have been limited by the technical difficulties of scaling from the human brain (950–1800 cm^3^) down to the mouse brain (0.32–0.52 cm^3^) [7–9]. Routine rodent brain connectomics is challenging because of the simultaneous demands of 1) reducing voxel volume by a factor of 3000, 2) providing sufficient diffusion weighting, 3) sampling an adequate number of gradient angles, 4) maintaining good signal-to-noise ratios, and 5) maintaining a reasonable scan time of less than 24 h per brain. All these requirements must be viewed in light of recent papers that highlight the problems of false negative tracts in structural diffusion connectomics [10, 11]. A single 1.25 mm^3^ voxel, as used in the human connectome project, can encompass on the order of 100,000 neurons and their processes [12]. Errors in tractography that result from crossing and merging fibers can result in substantial false positive connections. We have addressed many of these technical challenges by developing improved MRI protocols and unique hardware and encoding methods [13, 14]. We have validated and streamlined methods [9, 13] that enable the required high throughput for quantitative genetic studies of large murine cohorts [13, 14].

The major goal of this study was to systematically estimate heritabilities for many brain structures and parameters of a small but diverse cohort of mice. We measured volumes, scalar DTI metrics, and connectomes for each of 166 regions of interest (ROI), bilaterally, including both grey matter and fiber tracts. Most of these ROI are subcortical structures and tracts, many of which are comparatively difficult to quantify in humans. In total, we estimated volumes and DTI parameters for cortical regions, subcortical forebrain nuclei, midbrain regions, hindbrain structures, cerebellar regions, and white matter tracts. Finally, we estimated the maximum residual coefficient of errors with the conservative assumption that the exclusive cause of right-left differences was technical instead of due to genuine structural asymmetry [15]. Error terms were almost always under 5%, even for the smallest subcortical regions and tracts.

We addressed the following questions:

- What are the heritabilities of regional volumes and the different types of scalar metrics generated by DTI? Are connectome patterns heritable? Are there notable sex differences in heritability?
- To what extent are heritabilities compromised by errors of measurements, or by the size and shape of structures and tracts.
- How do differences in heritabilities and coefficient of variation map onto conventional neuroanatomical divisions and systems?

## RESULTS

### Whole brain population atlases provide statistical measures of heritability

Differences in brain size and shape, regional volumes, and even fiber tracts were notable between strains but only rarely notable within strains or between sexes (Figure 1, Figure 2, and Figure S1, S2). Our analysis highlights not only the more obvious morphological differences but also significant quantitative differences in the white matter anisotropy. For example, in the color fractional anisotropy image (ColorFA), the corpus callosum was much thicker and the saturation higher for the B6 line, which indicated higher anisotropy compared with D2 and CAST. At the extreme, the midline connection of the corpus was entirely absent in BTBR. In Figure 1, even the finest details of shape parameters, for example the entry of olfactory nerve fibers into the olfactory bulb, were highly distinctive from one strain to another, but not between males and females (compare top and bottom male and female averaged brain images). Although we have not explored shape parameters in this work, they may provide additional differential phenotypes. We limited this study to an analysis of variability and heritability of 1) regional volumes; 2) scalar DTI metrics, e.g. fractional anisotropy (FA), radial diffusivity (RD), axial diffusivity (AD), and mean diffusivity (MD) and 3) patterns of fiber connections.

**Figure 1:**
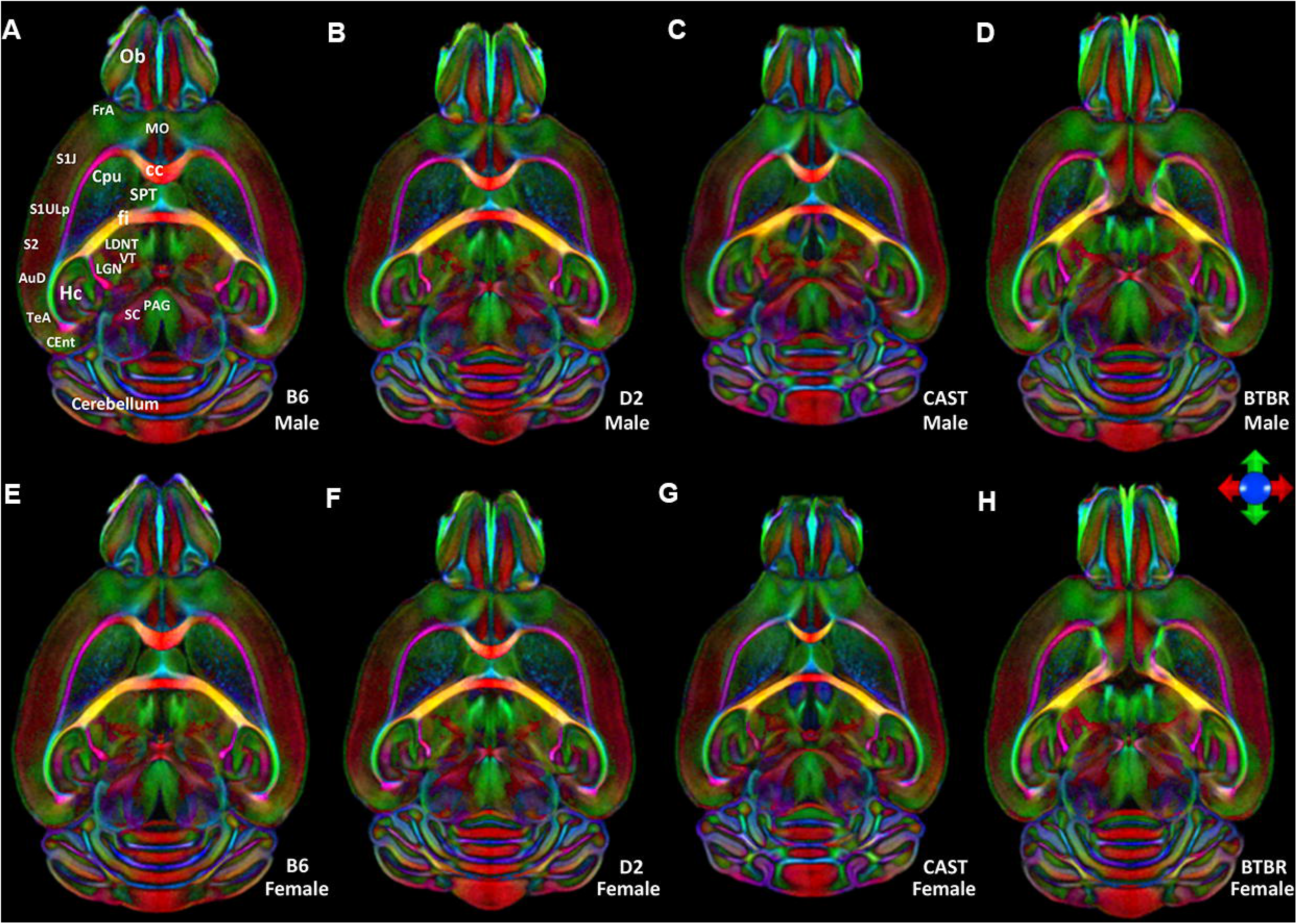
Horizontal color FA images from the average male (top) and average female bottom). Images are color-coded averages of four cases per sex that highlight orientation of fiber tracts (the principal eigenvector). The icon shows the key for encoding the eigenvectors: green-anterior/posterior; blue-dorsal/ventral; red-left/right. **A** and **E)** B6, **B** and **F)** D2, **C** and **G)** CAST, **D and H)** BTBR. Ob: olfactory bulb; FrA: Frontal association Cortex; MO: Medial Orbital Cortex; S1J,S1ULP;: Primary Somatosensory Cortex;S2: secondary somatosensory cortex; AuD: secondary auditory cortex; TeA: Temporal association cortex; CEnt: caudomedial entorhinal cortex; HC: hippocampus; SC: superior colliculus; PAG: Periaquaductal grey; LGN: lateral geniculate nucleus; VT: ventral thalamic nuclei; LDNT: latero dorsal nucleus of thalamus; fi: Fimbria; cc: corpus callosum; SPT: Septum; Cpu: striatum

**Figure 2:**
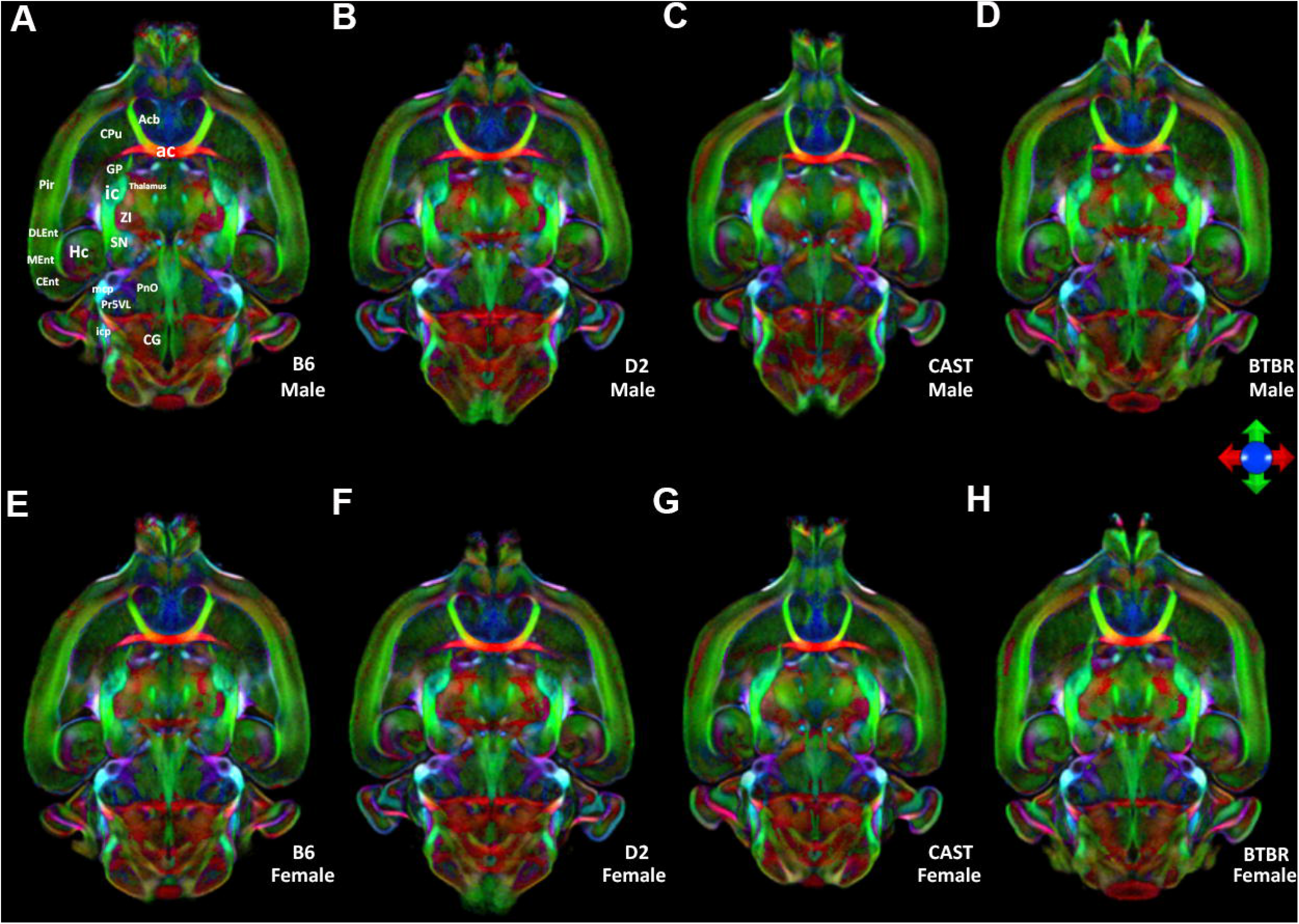
Horizontal color FA images from the average male (top) and average female bottom) at a second level. Images are color-coded averages of four cases per sex that highlight orientation of fiber tracts (the principal eigenvector). The icon shows the key for encoding the eigen vectors: green-anterior/posterior; blue-dorsal/ventral; red-left/right. **A** and **E)** B6, **B** and **F)** D2, **C** and **G)** CAST, **D and H)** BTBR. Acb: accumbens nucleus; Cpu: striatum; ac: anterior commissure; GP: globus pallidus; ic: internal capsule; Pir: piriform cortex; ZI: zona incerta; Hc: Hippocampus; DLEnt: dorsal intermediate entorhinal; MEnt: medial entorhinal cortex; Cent: caudomedial entorhinal cortex; mcp: middle cerebellar peduncle; PnO: pontine reticular nu, oral part; Pr5VL: principal s trigeminal, ventrol; icp: inferior cerebellar peduncle; CG: central gray.

The ColorFA is but one of ten different scalar images derived from the diffusion processing pipeline. Each image highlights different anatomy in a complementary fashion. Supplemental Figure S3 shows a comparison of the alternative contrasts. For example, the diffusion weighted (DWI) and FA were particularly useful in driving the label registration (See Star Methods).

### ROI volumes were heritable

For the most basic measurement, whole brain volume, our results are largely consistent with previous literature in volumes for male/female, respectively (in mm^3^): B6 463 ± 11.3 vs 479 ± 3.7; D2 415 ± 8.8 vs 403 ± 4.5; CAST 363 ± 11.98 vs 395 ± 4.6; BTBR 414 ± 7.8 vs 487 ± 3.9 (Figure S4). However, the BTBR brain volume we measured was smaller than B6, not larger as has been reported [http://gn2.genenetwork.org/show_trait?trait_id=49909&dataset=MDPPublish]. There were highly significant strain differences (F = 146.262, df = 3, *p* = 1.49e-15). mm^3^

### Sex differences were minor

We next wanted to determine whether sex was an important driver of individual differences. For every scalar parameter (volume, FA, RD, AD, MD), ANOVA comparisons were carried out including strain, sex, and their interaction.

There was no significant sex difference in total brain volume for male vs female (445 ± 20 SE versus 442 ± 18 SE). For volume of individual ROIs, 34 regions had a significant (B-H q ≤ 0.05) sex effect, 328 had a significant strain effect, and there was no significant strain by sex effect. The region with the most significant sex difference was the bed nucleus of the stria terminalis (BNST; right male = 0.0599 ± 0.0022, female = 0.0551 + 0.0032, p = 0.00002; left male 0.0599 + 0.0026, female 0.0556 + 0.00277, p = 0.0000780). This region is sexually dimorphic in humans [16]. Heritability estimates for volume were significantly correlated between sexes (r = 0.75).

In the scalar DTI metrics, there were fewer sex effects than for volume, and heritability estimates in each sex correlated well. For FA, two regions had a significant (B-H q ≤ 0.05) sex effect, 298 regions had a significant strain effect, and none had a strain by sex effect. There were no significant sex or sex by strain effects for RA, but 168 regions had a significant strain effect (B-H q ≤ 0.05). For MD, there were no significant sex or sex by strain effects, but 118 regions had a significant strain effect (B-H q ≤ 0.05). There were no significant sex or sex by strain effects for AD, but 109 regions had a significant strain effect (B-H q ≤ 0.05). These results showed that sex effects were relatively minor compared with strain effects, especially for DTI metrics.

### Significant strain differences in most of the scalar measures

The data in Figure 3 show that there were significant qualitative strain effects for all the scalar measures, but we wanted to quantify the extent to which strains and cases differed for any measurement. To fulfill this aim, we carried out ANOVA comparisons for each ROI and each measurement, using Tukey’s posthoc test to determine which strains were different. For volumetric data, 327 of 332 regions and tracts differed significantly (Benjamini-Hochberg corrected *q* < 0.05; Figure 3A). The five regions that did not differ were the left ventral tegmental area, right ventral orbital cortex, left dorsal acoustic stria, left lateral orbital cortex and right lateral orbital cortex. The differences in such a large number of regions confirmed our initial assertion, namely that BTBR and CAST would be most different from the ‘normal’ lab strains B6 and DBA2, but also that most brain regions would be significantly different between these two ‘normal’ strains. We observed similar results for the other measures: For FA, 298 regions had a significant ANOVA (Benjamini-Hochberg corrected p < 0.05), MD 128 regions, AD 117 regions, and RD 175 regions (Figure 3b-e).

**Figure 3:**
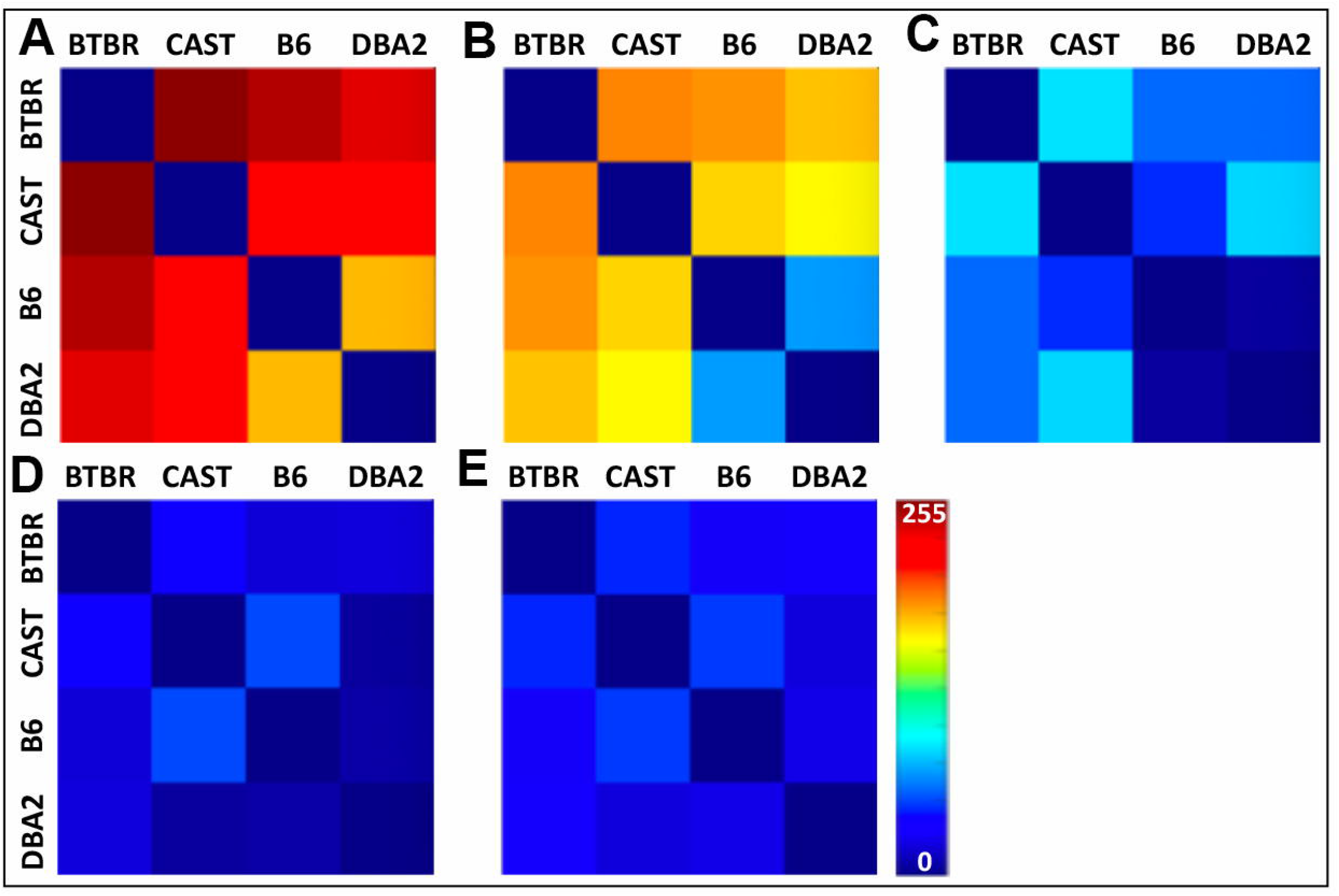
Heatmaps demonstrate the differences in volume and four of the scalar DTI metrics between strains. ANOVAs were performed to determine the number of regions of interest (from a total of 166×2) for which there were significant strain effects. A) Volume; FA; C) RD; D) MD; E) AD;

### Significant strain differences are heritable

For most brain regions, there was a high heritability (*h_2_*) of volume. The FA had a high *h_2_* for a more limited number of regions. In general, MD, AD, and RD had low *h_2_* (Figure 4).

**Figure 4:**
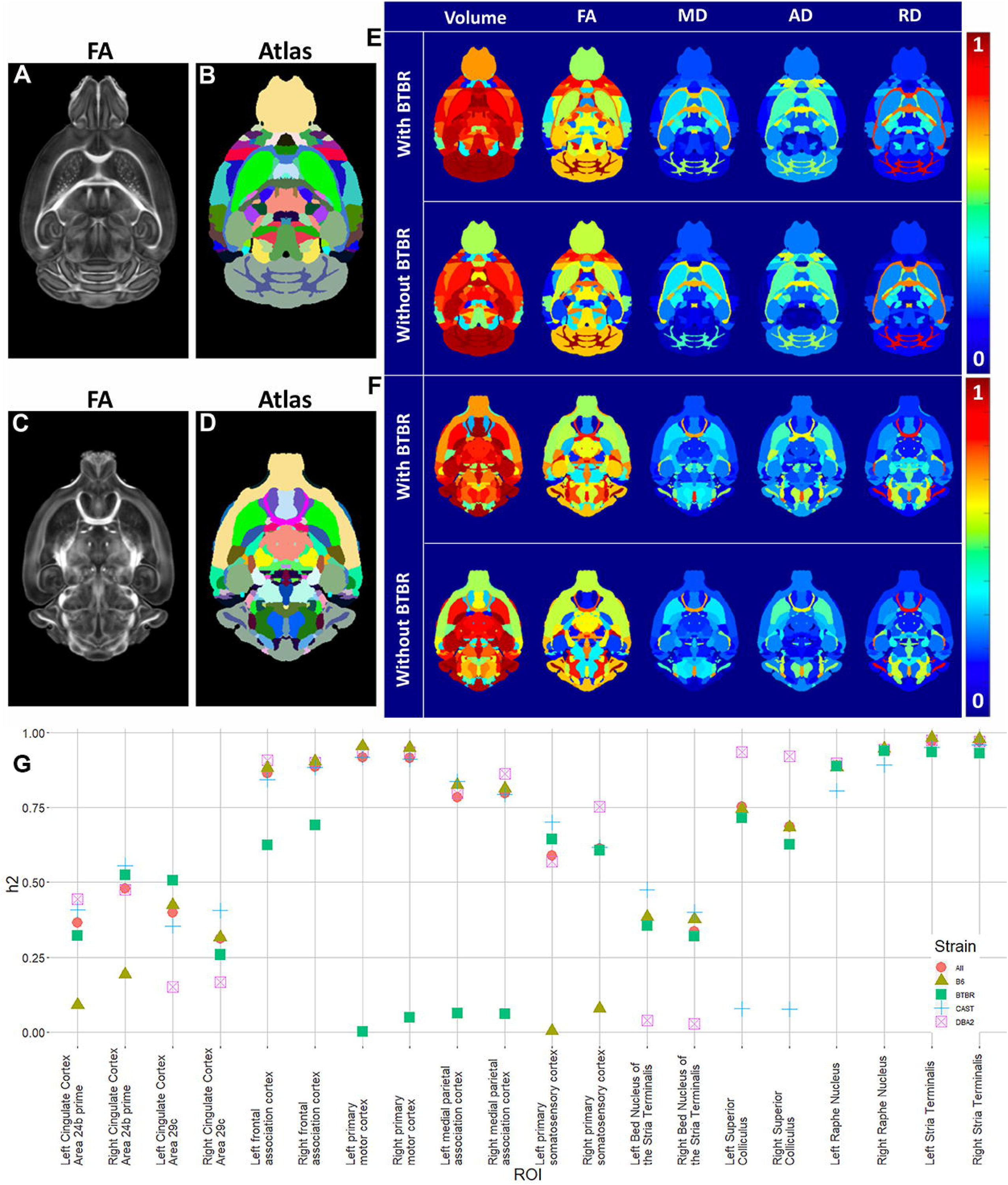
Multiple Biomarkers are Heritable. A) and C) Representative dorsal slices from the FA images of the B6 and B) and D) the reference regions of interest. E) and F) Heritability (h_2_) is shown for volume, fractional anisotropy (FA), mean diffusivity (MD), radial diffusivity (RD), and axial diffusivity (RD) for these same two reference slices. G) Heritability of the volume of 20 brain regions, across both hemispheres, dropping a strain (C57BL/6J = B6, DBA/2J = DB, BTBR = BT, CAST = CA, None dropped = No). ROIs are the hemesphere (left or right), our ROI number, and the region name

Often, a single outlier strain will drive these high heritabilities. To demonstrate this effect, we examined the heritability of the volumes of ten regions of interest, comparing estimates from the left and right hemisphere, recalculated when leaving one strain out (Figure 4G). The results demonstrate three important points. First, for many brain regions, one strain drove heritability because it was an outlier. Second, although BTBR and CAST were frequently outliers, there were also regions of DBA/2J or the ‘normal’ C57BL/6J that were outliers. Third, as expected, heritability was broadly the same across both hemispheres. The complete Table (S1) for this analysis is available in our on-line supplement.

The heritabilities for all measures in Figure 4E were not strongly dependent on their values, i.e. heritability was not necessarily higher for large values. This independence suggested that we could measure low values accurately; if low values had higher variability, they would have had lower heritability. Figure S5 shows the h_2_ ranking as ordered by ROI in a comparison of the four strains; Table S1 lists the top 20 regions from this rank-ordered list for each of the scalar metrics. Cerebellar white matter FA and FA from multiple regions of the cortex were highly heritable. Several regions were highlighted in both volume and diffusion changes, e.g., cingulate cortex- volume and FA; fimbria-volume, RD, and AD; lateral lemniscus- volume, FA and AD. Multiple regions had correlated heritability in AD, RD, and FA.

Figure 5 illustrates the high heritability (*h_2_*) for the volume of many white matter and cortical regions. A number of regions had high heritabilities of FA, and MD, but the heritability of AD, and RD was generally low. To determine whether certain brain systems have higher heritability than others, we calculated the average heritability for the volume in five broad regions: forebrain, midbrain, hindbrain, white matter tracts, and ventricular. An ANOVA test showed highly significant differences between these brain parts (df = 4, F = 6.435, p = 5.39e-05). Tukey’s posthoc suggested that the forebrain (mean h_2_ = 0.53, n = 180) drove these differences, with significant differences from hindbrain (mean h_2_ = 0.64, n = 62, p = 0.005879) and white matter tracts (mean h_2_ =0.67, n=64, p=0.0002724).

**Figure 5:**
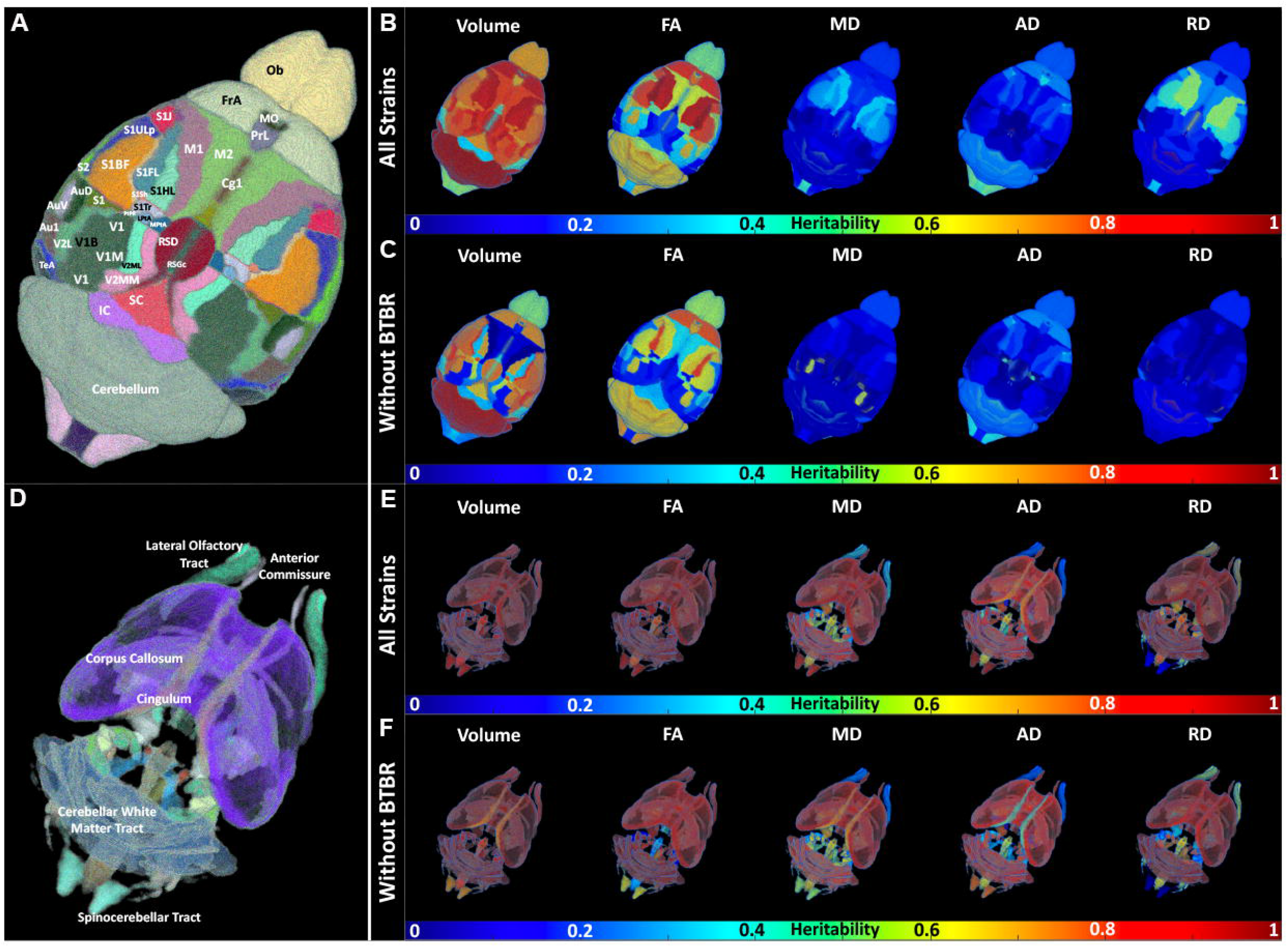
Spatial distribution of heritability in cortex and white matter regions with and without BTBR strain. Heritability (h_2_) is shown for volume, fractional anisotropy (FA), mean diffusivity (MD), radial diffusivity (RD), and axial diffusivity (RD).

We next wanted to determine whether one strain drove these differences, in a manner illustrated by Figure 4G. Table 1 shows the significance of the difference between mean Tukeys post-hoc, with all strains, or dropping each strain. The difference between white matter tracts and forebrain was significant in all comparisons, suggesting it is not dependent upon one strain. However, the CAST strain appeared to drive the difference between forebrain and hindbrain, suggesting that CAST was an outlier for heritability of this system.

**Table 1:** Tukey comparisons of mean heritability.

|  | All strains | D2 | B6 | CAST | BTBR |
| --- | --- | --- | --- | --- | --- |
| Midbrain - Forebrain | 0.828 | 0.447 | 0.999 | 0.823 | 0.845 |
| Hindbrain - Forebrain | 0.006 | 0.004 | 0.072 | 0.999 | 0.0007 |
| White matter tracts - Forebrain | 0.0003 | 0.0003 | 0.003 | 0.015 | 0.006 |
| Ventricular system - Forebrain | 0.317 | 0.097 | 0.757 | 0.204 | 0.812 |
| Hindbrain - Midbrain | 0.855 | 0.989 | 0.505 | 0.912 | 0.653 |
| White matter tracts - Midbrain | 0.603 | 0.9187 | 0.195 | 0.927 | 0.851 |
| Ventricular system - Midbrain | 0.773 | 0.653 | 0.798 | 0.647 | 0.991 |
| White matter tracts - Hindbrain | 0.971 | 0.986 | 0.925 | 0.117 | 0.986 |
| Ventricular system - Hindbrain | 0.964 | 0.754 | 0.999 | 0.265 | 0.998 |
| Ventricular system - White_matter_tracts | 0.994 | 0.863 | 0.999 | 0.852 | 0.999 |

This table shows the significance of Tukey posthoc for comparisons of mean heritability across five brain systems, for all strains, and drops each strain individually. D2 = DBA/2J, B6 = C57BL/6J, CAST = CAST/EiJ, BTBR = BTBR T+ *Itpr3<sup>fl</sup>/J*,

### Sex and hemisphere dependent coefficient of variation

The coefficient of variation [17] of individual ROI volume between left and right hemisphere for four strains were the following: D2: 0.0158±0.0168; B6: 0.0157±0.0199; CAST: 0.0156±0.0163; BTBR: 0.0140±0.0152. Figure S6 shows the CV of individual ROI as a function of the ROI volume. For most of the ROI in all strains, the CV was <0.05. The CVs between male and female consistently showed low values: D2: 0.0373±0.0301, B6: 0.0307±0.0265, CAST: 0.0366±0.0309, BTBR: 0.0282±0.0276;

### Connectomes for each strain

Figure 6A shows the whole brain connectome for all four strains: a) B6; b) D2; c) CAST; d) BTBR. The connection strength is shown with a log10 color scale that demonstrated an enormous range in connection within a single brain and across the strains. Each graph includes four sub sections: upper left- connections within left hemisphere, upper right- connections from left hemisphere to right, lower left- right hemisphere to left, and lower right- connections within right hemisphere. Visual comparisons reveal differences between the BTBR and the other strains. Reduction between the hemispheres is evident when one examines the upper right and lower left section of each quadrant. There are some intriguing differences between the more normal strains and the BTBR. The upper right-hand part of each quadrant shows connectivity between the isocortex and midbrain, hindbrain and white matter. The edge strengths in all these regions are considerably reduced in the BTBR. For example, the black arrows highlight the corpus callosum of the B6 and BTBR. Expanded plots of these nodes show the strength of connection between left and right hemisphere. The log color scale highlights the reduced connection between left and right hemisphere for the BTBR.

**Figure 6A.**
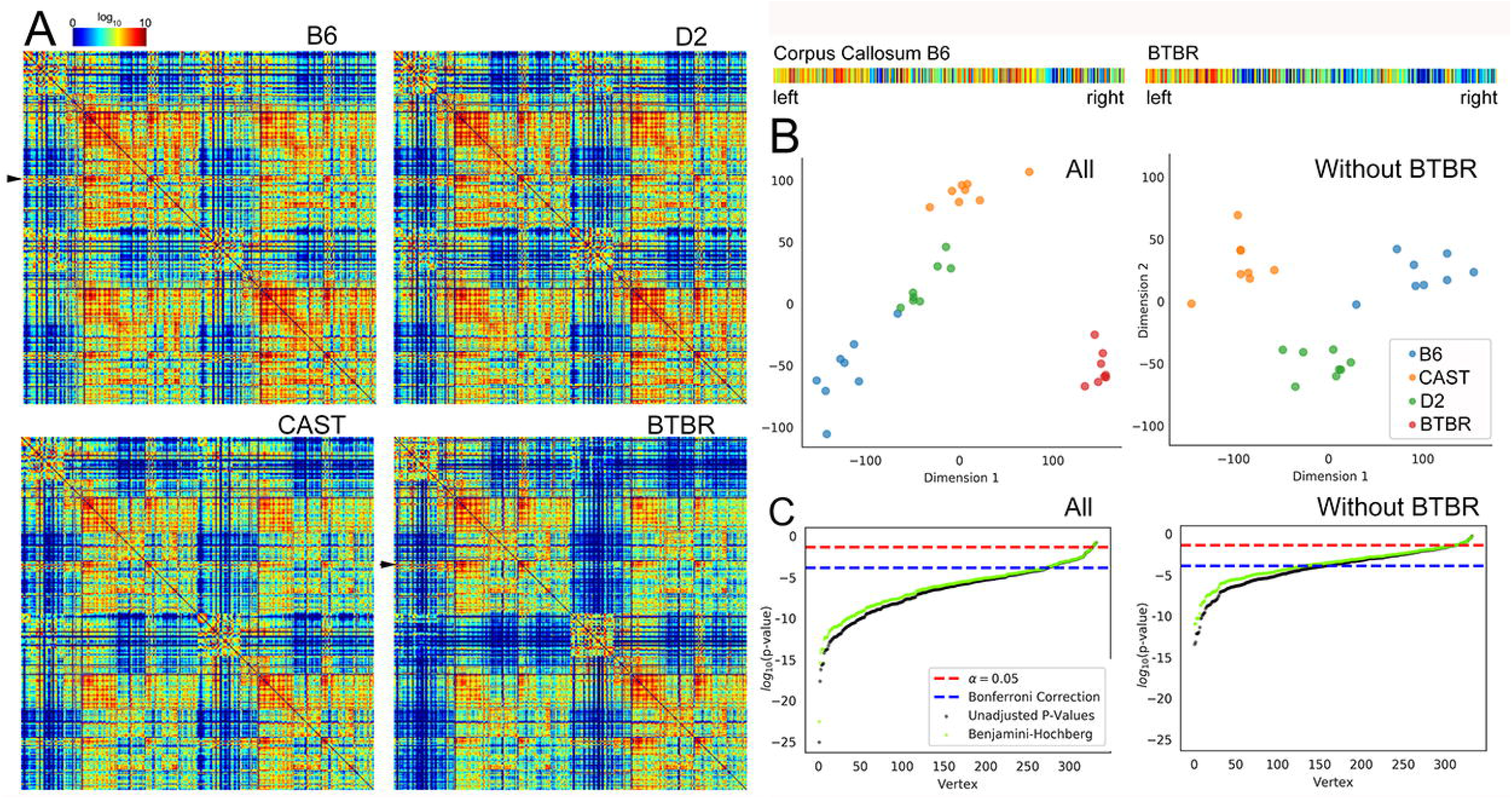
Whole brain connectome for each strain. The connectomes (n=8) of a) B6; b) D2, and c) CAST strains were similar. The average connectome of the BTBR was significantly different with the reduced connectivity evident from the increased number of holes (dark blue entries). The seed nodes [22] and targets (columns) are ordered according to the developmental origins. The arrows identify the corpus callosum (seed) which was extracted and magnified for comparison between B6 and BTBR. **Figure 6B: Omnibus Embedding Improves the Confidence in Comparisons**. The scatter plot shows clustering when the BTBR was included. When the BTBR was excluded, the clustering also shows statistical difference in the three similar strains. **Figure 6C:** Plot of log (p-value) from MANOVA comparisons in vertices across strain vs rank-ordered vertex shows roughly 280 vertices that were statistically different when all four strains were compared and 150 vertices that were different when the BTBR was excluded.

Although subjective comparisons of connectomes in Figure 6A suggest regions where there were differences, statistical comparisons were challenging. First, false discovery was a confound because of the size of the array (332×332) [35]. Second, the heritability calculations were essentially test statistics, whose sensitivity and specificity depended on the degree to which the data satisfied certain assumptions. Although those assumptions are often appropriate, for these data, exploratory analysis demonstrated that the assumptions were not valid. Nonparametric methods for quantifying heritability failed to rectify this situation. Figure S7 demonstrates that no edges on their own passed the typical 0.05 level of significance once we conservatively corrected for multiple comparisons. The problem stemmed from the fact that existing methods for quantifying these differences were not adequate. Therefore, we devised a new method for statistically determining heritability of connectomes. Instead of treating each edge as an independent object, we first assessed whether the connectome, as a whole, was heritable.

Indeed, as demonstrated by Figure 6B, the connectomes of a given strain tended to be more similar to one another than they were to the connectomes of other strains. Using classical multidimensional scaling to embed each connectome into just two dimensions, we found that the connectomes tended to form strain-specific groups. This connectome-wide group effect was preserved when we included and excluded the BTBR strain. With these results, the natural next question was “Which brain regions confer the resulting heritability?”

To answer this question, we leveraged recent developments in the statistical analysis of populations of graphs on a shared vertex set. This enabled us to find a joint low-dimensional representation of each node in each connectome, based on its connectivity profile, that is, its set of connections. In this way, we could use standard methods for subsequent inference. Particularly, we posited a multivariate, non-parametric k-sample test hypothesis: For each node, was the low-dimensional representation for a given strain sampled from the same distribution as the other strains? Figure 6C shows that about 280 nodes, of the 322 possible nodes, were significantly different across all four strains, after a stringent correction for false positives. The same analysis for the three control groups (excluding the BTBR) demonstrated that there were about 150 nodes that were significantly different. Moreover, there were multiple pairs of nodes for which both hemispheres were significantly different. Table S2 presents the top twenty nodes.

Figure 7 shows the tractograms for four of the nodes in which the p values were the most significant. The corpus callosum was profoundly different in the BTBR, and it was the most highly ranked node when we compared all four strains. The termination at the mid line is obvious in Figure 7b. The corpus callosum was also highly heritable when we excluded the BTBR. And this was consistent with the anatomical observations from Figure 1 in both the morphology and intensity in the color FA. The substantia nigra, internal capsule, and fimbria were the next three most highly ranked nodes in the comparison of all four strains. The internal capsule and substantia nigra were also in the top four nodes when we compared the three similar strains, thus, these regions may be highly heritable across many strains. Examination of the tractograms affirmed the statistical findings. Tracts across the midline were substantially reduced for the BTBR in the substantia nigra and internal capsule and clearly less dense in the contralateral side for the fimbria compared with the other three strains.

**Figure 7:**
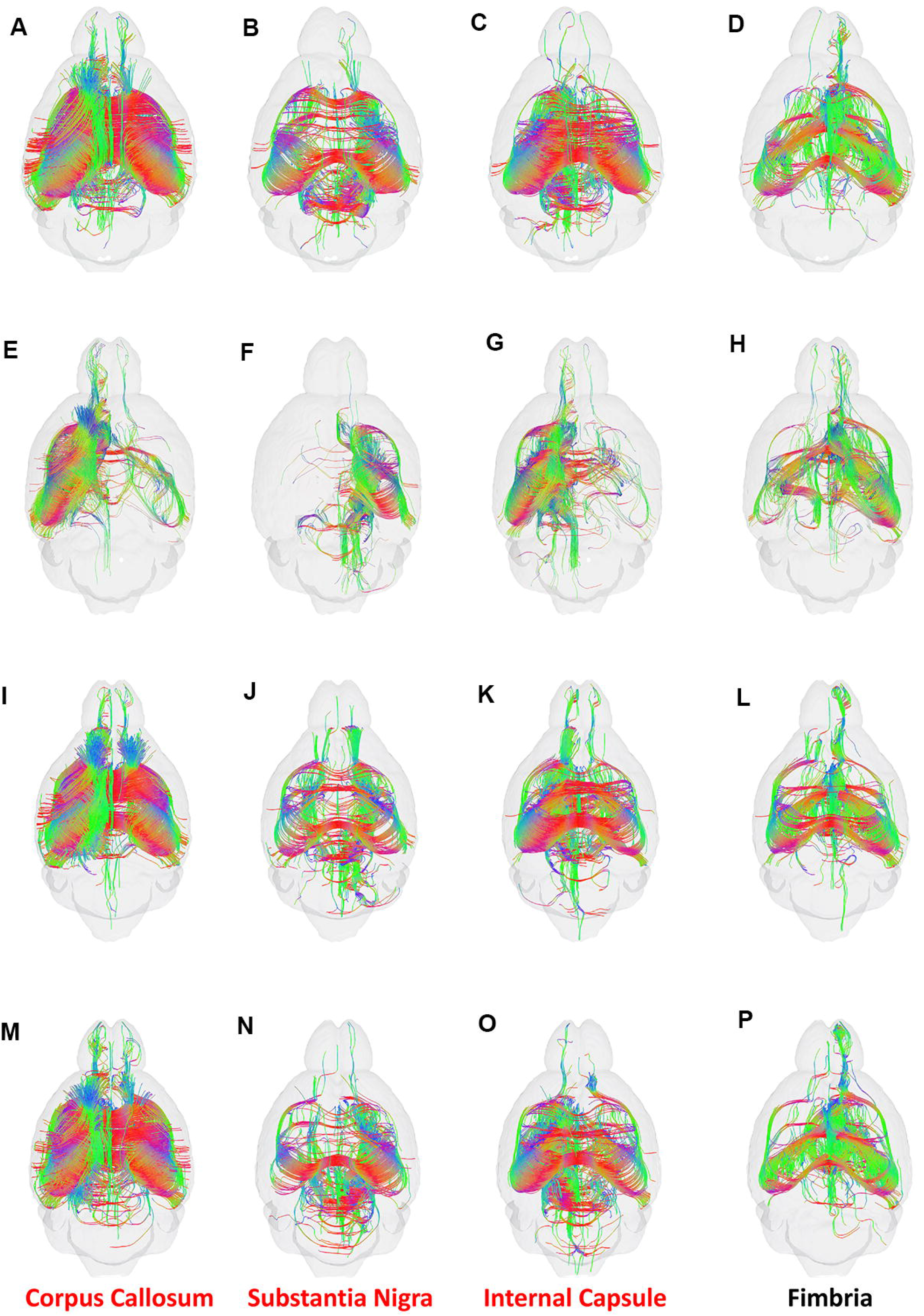
Tractograms for the top 4 nodes. derived from the omni MANOVA analysis including all four strain. A),E),I)M); B6, B),F),J),N; BTBR, C),G),K),O); CAST, D),H),L),P); D2. The Corpus callosum, Substantia Nigra and internal capsule were also statistically different when the BTBR was excluded.

## DISCUSSION

We have used state-of-the-art MRI workflows to study heritabilities in the mouse brain at three different levels: 1) volumetrically-defined brain structures, including more than 100 regions of interest—both nuclei, cortical regions, and fiber tracts, 2) scalar DTI parameters that were measures of tissue cytoarchitecture, and 3) variation in DTI-defined connectomes. For this initial work, we used four fully isogenic inbred strains of mice that differed greatly in genome sequence-—from 5 to 15 million variants distinguished any pair of strains.

The most important findings are the following:

### Microscopic MRI/DTI provides quantitative neuroanatomical phenotypes

Brain connectomes can vary over an enormous range of scale, over species, and between technologies. The electron microscopy connectome of *Drosophila* that was acquired at nanometer resolution was constructed at voxels on the order of 10^-15^ mm^3^ [18]. Clinical MRI acquires data with voxels on the order of 2 mm^3^ [12]. The mouse brain data in these studies were acquired with voxels < 10^-5^ mm^3^. Tract tracing in an EM image relies on different physical phenomena compared with a retroviral tracer which, in turn, is a considerably different tracing mechanism compared with diffusion. Thomas et al. [10] and Maier-Hein et al. [11] have explored the specificity and sensitivity of the diffusion based (human) connectome and concluded that frequently there can be more false positive connections than true positive connections. One explanation for a preponderance of false positives is that merging or crossing fibers can cause errors in tracking. This observation has driven us to push the spatial/angular resolution (in rodents) by nearly 5 orders of magnitude beyond that of clinical scans to limit the consequences of volume averaging [9]. The data reported here are, to the best of our knowledge, the highest (spatial/angular) resolution available of genetically linked diffusion connectivity metrics in the mouse brain. And we have carefully catalogued the limits of sensitivity and specificity relative to retroviral tracers [9, 13]. Several investigators have used in vivo MRI to investigate structural connectivity in genetically engineered mice. Practical problems of life support limit the resolution in these studies to ~100×100×500 um with ~30 angular samples [19, 20]. Scans of post mortem specimens enable longer, higher resolution studies. In 2002, Zhang et al. reported some of the first DTI images of the post mortem mouse brain [21], and, more recently, they have used the method to understand the function of Bcl-x in brain development [22]. Subsequent studies have extended spatial resolution and applied the method to unraveling morphologic changes in genetic models of neurologic disease [23, 24]. These studies relied on the scalar metrics (AD, RD, FA etc.), but they did not include any connectome. Generating the connectome adds a significant complication because one must acquire a “sufficient” number of angular samples to enable resolution of crossing and merging fibers. The term “sufficient” has been controversial. Calamante et al. have developed higher resolution tract density methods for connectomics in post mortem specimens at 100 um^3^ with 30 angular samples [25]. To enable comparison across the range of applications (human, mouse, in vivo, post mortem), we proposed the resolution index (RI), i.e., the product of the spatial and angular resolution [14]. RI for the human connectome was 128 [12]. RI for Calamante’s study was 30,000 [25]. The RI for this work is >500,000.

### Volume, FA, AD, RD, and Connectomes are Heritable

Virtually all the scalar metrics we measured were heritable at some level in some regions. Multiple investigators have demonstrated heritability of volumes in specific sub regions of the brain in both human and rodent studies [26–29]. Other recent studies have detected significant global heritability of FA, AD and RD in human populations [6, 30]. And studies have demonstrated heritability of specific regions of connectivity, in human [31, 32]. However, the work described here is the first to bring together a robust study of heritability of all these metrics (volume, FA, AD, RD, connection strength) in the mouse. We have demonstrated region specific heritability of volume, FA, AD, RD and connectivity.

We evaluated three strains (B6, D2, CAST) for which we expected neuroanatomical and connectome differences to be comparatively modest after correcting for global differences in brain volume. We expected a fourth strain (BTBR) to display major disruptions of interhemispheric connections. Working with isogenic animals (n = 8/strain) has enabled us to measure the within-strain variability of all traits with good precision, but with the caveat the we have sampled only a small number of genomes. Nonetheless, we could test the methods in detection of major differences (e.g. the BTBR) when the anatomy was so different that one needed to be concerned about artifacts from label registration and other experimental errors in the processing pipeline.

There was a remarkably robust collection of heritable phenotypes. Volume was highly heritable across the entire range of the structures included. Previously, we demonstrated that genotype accounted for ~ 60% of the variability of volume differences in 33 regions of the brains of a cohort of C57BL/6 mice and ten different strains from the BXD family [33]. The average coefficient of variation of volume for the 33 structures across 11 strains was 15.2 ± 5.8%. The new studies described here provide the same data for 166 ROIs (in both hemispheres) with substantially improved precision. As shown in Figure S6, the coefficient of variation within a strain for most of the regions was <5%. The CV increased for smaller structures, i.e., structures with volumes <0.1 mm^3^ [34].

There were only nine ROIs (1.4%) with volumes >0.1 mm^3^ with CV>5%. The heritability was not strongly dependent on the absolute volume of the structure. More than 150 structures per hemisphere were highly heritable (>0.8). Previous GWAS analysis of aggregated data from fifty other studies demonstrated heritability of volume in subcortical regions [5, 35]. The VETSA study compared 1,237 twins by focusing on heritability of cortical thickness [36]. The ENIGMA consortium expanded on the heritability of subcortical structures by comparing lateral symmetry using 15,847 MRI scans from 52 different data sets [37]. Elliott et al. used data from the UK Biobank to measure heritability across morphometric, diffusion and functional data in 8,428 individuals [4].

Kochunov et al. demonstrated that whole brain FA and RA had strong genetic dependence, whereas the genetic link to AD was not significant [6]. A study of twins by Gustavson et al. revealed that all of the regions they measured were heritable and FA and MD accounted for roughly half the heritability in the tract subdivision [38]. Recent studies from the UK Biobank have explored a number of image-derived phenotypes demonstrating regional heritability of FA, AD and RA [4]. Our work shows similar results with FA having the highest heritability of the scalar diffusion metrics. In the full group (including BTBR), the FA was strongly heritable (>0.8) in ~130 of 166 regions of interest. Surprisingly, in the smaller group of more similar strains, heritability was still >0.8 in more than 100 of the ROIs. AD and RD were not linked as strongly. But heritability was still >0.8 in 50 ROIs for RA and in ~30 ROIs for AD.

Finally, the Human Connectome Project has generated multiple studies of heritability of specific structural circuits in the brain. Figure 5 shows, again, a remarkably rich set of vertices (~280) with strong heritability across both hemispheres. As with the scalar metrics (Figure 2), the evidence of genetic dependence in the connection strength was still present when we removed the BTBR strain from the analysis. A statistically robust comparison of connectomes is challenging because of false discovery. Trends that seem apparent in the color-coded adjacency matrices are influenced by the brain’s ability to integrate the patterns. Creation of a connectivity profile for each node exploits this effect of integration and yields a more statistically robust method to define the nodes that have the most difference across the strains.

## CONCLUSION

The brain is the most complicated organ in the body. Understanding its genetic underpinnings starts with an understanding of brain structure. But the structure is fractal with collections of neurons linked through axons, collections of axons linked to local circuits, and circuits classified as short or long range. No single view or scale provides a complete picture, and no single method is sufficient to provide the full picture of the brain. Unwinding the genetic links to brain structure provides crucial knowledge to understanding a great many diseases and disorders (such as autism, dementia and Alzheimer’s disease). Sorting through the extraordinary complexity of data can quickly become overwhelming. Rank ordering the scalar metrics and the most critical nodes in the connectome brings specific circuits in focus. Sophisticated network analysis can then be used to construct hypotheses that connect structural changes to behavior. The methods shown here enable one to span the scale from meso to micro over the entire brain. When coupled to the power provided through isogenic colonies, our method promises a statistically robust approach to linking connectome to genome.

## ACKNOWLEDGEMENTS

We are grateful to Gary Cofer, James Cook, and Lucy Upchurch for invaluable technical assistance. We thank Tatiana Johnson for special care in manuscript preparation and submission.

## AUTHOR CONTRIBUTION

NW acquired the DTI data, performed much of the data reduction, generated most of the figures, and assisted in writing the manuscript. RJA developed the annotation set, performed the label registrations, and assisted in preparing figures. DGA assisted in the heritability analysis, generation of figures, and writing the manuscript. VG, YP, CEP, and JTV developed the statistical approaches and assisted in preparation of figures. JYV assisted in manuscript preparation. YQ performed tissue preparation and managed data provenance. RWW and GAJ designed the experiment and wrote the manuscript.

## STAR METHODS

Detailed methods are provided in the online version of this paper and include the following:

- LEAD CONTACT AND DATA AVAILABILITY
- EXPERIMENTAL MODEL AND SUBJECT DETAILS
- METHOD DETAILS

- Image Acquisition
- Image Registration
- Connectome Generation
- DATA AVAILABILITY

## LEAD CONTACT AND DATA AVAILABILITY

Further information and requests for data should be directed to the Lead Contact, G. Allan Johnson

## EXPERIMENTAL MODEL AND SUBJECT DETAILS

The Duke Institutional Animal Care and Use Committee (IACUCC) approved all the study protocols. We compared four mouse strains.

1. C57BL/6J (B6)—often referred to as “the mouse”—is the references strain for murine genomics and neuroscience. The B6 is the maternal parent of the large BXD family of strains (*n* of 150 progeny strains; [39]). B6 is a member of the *Mus musculus domesticus* subspecies, and it has a mean brain weight of 479 ± 3.7 mg (*n* = 259, all brain data generated in our laboratory using relatively uniform methods, [40] and www.mbl.org/references/BrainWeightOfStrainofMice.xlsx, and GeneNetwork trait 49909).
2. DBA/2J (D2)—the oldest fully inbred strain—is the parent of the BXD family, and, like B6, it has also been sequenced [41]. D2 is also a *Mus musculus domesticus*, and it has a mean brain weight of 403 ± 4.5 mg (n = 154).
3. CAST/EiJ [27] is a fully inbred strain, derived from the Southeast Asian *Mus musculus castaneus* subspecies. This strain breeds well with other subspecies, and, although it is somewhat smaller, it has a mean brain weight of 395 ± 4.6 mg m (*n* = 23).
4. BTBR T+ *Itpr3tf*/J (BTBR) is a common inbred strain with well-known behavioral and neuroanatomical abnormalities including a highly penetrant callosal defect [42] [43] [44]. BTBR is a *Mus musculus domesticus* with a brain weight of 487 ± 3.9 mg, close to that of B6 (*n* = 51)

The first three strains have comparatively normal CNS structure and connections, whereas BTBR is an acallosal strain [42] [43] [44]. There is a 20% range of difference in mean brain volumes among these strains; B6 and BTBR have relatively large brains, whereas D2 and CAST have smaller brains [39]. Thus, we corrected all heritability estimates to eliminate effects of different overall brain volume.

Four males and females for each strain were purchased from The Jackson Laboratory. Brains were left in the cranium and actively stained via perfusion fixation with Prohance, a gadolinium contrast agent that reduces the spin lattice relaxation time (T1) [45]. Mandibles were removed to enable use of a smaller radiofrequency coil. Specimens were mounted in a 12-mm-diameter plastic cylinder. The cylinder was filled with fomblin, an inert fluorocarbon that reduces susceptibility artefacts. All animals were young adults at 90 days of age.

## METHOD DETAILS

### Image Acquisition

MR images were acquired on a 9.4T vertical bore magnet with Resonance Research gradients providing ~2000 mT/m maximum gradient. The system was controlled by an Agilent Direct Drive console. Specimens were mounted in a 12 mm diameter single sheet solenoid radiofrequency coil. Three-dimensional (3D) diffusion weighted images were acquired with a Stejskal Tanner rf refocused spin echo sequence with TR/TE of 100/12.7 ms and b values of 4000 s /mm^2^. Forty-six volumes were acquired, each with a different gradient angle along with five baseline (b0) images distributed throughout the four-dimensional (4D) acquisition. Sampling angles were uniformly distributed on the unit sphere. Compressed sensing was used with an acceleration of 4 X to reduce the acquisition time to 23.2 h/specimen [13]. The result was a 4D image array with isotropic spatial resolution of 45 um (voxel volume of 91 pl).

### Image Registration

The 4D array (256×256×420×51) was passed to a post-processing pipeline that first registered the five *b0* images together. Then each of the diffusion-weighted 3D volumes was registered to this average to correct for eddy current distortion. The registered 4D array was passed to DSI Studio which generated scalar images: Axial Diffusivity (AD), radial diffusivity (RD), mean diffusivity (MD), fractional anisotropy (FA), and color fractional anisotropy (ClrFA) using the DTI model [46]. Representative images (Supplement Figure S3) demonstrated the complementary anatomic detail provided by these scalars to assist in the label registration.

### Label Registration

A 3D label set was registered onto each 4D volume in that volume’s native reference frame, that is, the reference frame in which the data were acquired. A symmetric Waxholm Space (WHS) [47], i.e., an isotropic label set of 166 total ROIs in each half of the brain [9], was generated by reflecting the label set through the midline. The symmetrized label set minimizes bias in lateral comparisons [48]. The complete list of labels is available in the Supplemental Table S3. The labels were isotropic, i.e., they extended in all three planes at the same spatial resolution (45 μm) (Figure S8).

### Coefficient of Variation

Left and right labels were generated for each of the 32 specimens by creating a strain- and subject-specific spatial mapping to WHS. These maps are the source images (DWI, FA, AD, RD) referred to here as “WHS-Atlas”, including the label set of 166 regions per hemisphere. Figure S9 summarizes the path of the labels from the WHS Atlas to the individual specimen [49]. Because there were significant structural differences among the strains that might have pushed the registration pipeline beyond its capabilities, an intermediate hybrid average was generated for each strain. Averages were generated using a pipeline incorporating ANTS [50] with the following steps:

1. An automated skull stripping algorithm removed all the external tissue from the DWI and FA images;
2. A second stage of affine transform was applied, again using the DWI;
3. Six iterations of non-linear registration were performed using the diffeomorphic Symmetrized-Normalization algorithm as implemented in the Advanced Normalization Tools software [50, 51], and was driven by the FA contrast.

After each diffeomorphic warp, images were averaged to create a new template. An average of all the inverse warps was then applied incrementally to move the shape of this template towards the median shape of the group before being used as the target of the next iteration. The strain average for each strain was created from eight specimens in that strain. Merged strain averages were created for each strain/gender by using four B6/gender cases with the four data volumes of the strain/gender to which the labels were being mapped. Linear registrations used the Mattes similarity metric, whereas non-linear jobs used cross-correlation. For D2, the WHS-symmetric labels were warped using the appropriate transform chain and nearest neighbor interpolation such that each template had its own unedited label set. Before being propagated to the individual specimens, the template labels in the strain average were inspected and manually edited. It was necessary to refine the white matter of the cerebellum via thresholding of the template FA image. D2 and CAST required slight manual edits, primarily involving the boundary of the caudate-putamen, whereas significant edits were made to the BTBR template labels due to notable anatomical differences in the cortex and midline structures.

### Connectome Generation

Tracts were generated using the generalized Q-sampling (GQI) fiber tracking algorithm in DSI studio with a maximum of three fibers per pixel [52]. The propagation direction was calculated by applying trilinear interpolation on the fiber orientations provided from neighborhood voxels. The next point was then determined by moving 0.02 mm in the propagation direction. The propagation process was repeated until the tracking trajectory exceeded either a turning angle of greater than 45° or the anisotropy value of the current position was below a predetermined threshold. Five million fibers were generated with whole brain seeding for a reference B6 specimen. The number of fibers for all other specimens was normalized to each specimen volume. For whole brain tractography (white matter and gray matter), the FA threshold was 0.1 with a minimum fiber length of 0.5 mm.

Whole brain connectomes were generated for each specimen in DSI studio using the registered label set with 166 regions of interest on each hemisphere. The seed and target regions were ordered based on their developmental origins. A complete key for region number and its anatomical definition is included in Supplemental Table S3.

### Heritability Analysis

Heritability is the proportion of phenotypic variance that is explained by genetic differences. In fully homozygous strains, additive genetic variance is the main source of differences between strains, and our estimates can be considered close to what is typically called narrow sense heritability (*h_2_*). *h_2_* was estimated as the fraction of variance explained by strains in a simple ANOVA model.

We also calculated H2RIx̅ [53], the heritability of differences of strain means. This alternative estimate of heritability accounts for the number of within-strain replicates, and will always be higher than *h_2_*, when *n* >1. Formally, H2RIx̅ is defined as H2RIx̅ = V_a_/(V_a_ + V_e_/n), where V_a_ is the variance among strain averages (typically four strains in our study), V_e_ is the environmental variance (the average variance of the within-strain variances), and *n* is number of replicates within strain (eight in this study).

These two estimates have different uses. Heritability in human studies and conventional intercrosses are reported as *h_2_*, therefore, this value is best for comparisons to previous work. In contrast, H2RIx̅ takes into account the reduction of technical and environmental sources of variance achieved by resampling. Therefore, H2RIx̅ is most useful when estimating power of MRI studies that take advantage of replicable isogenic populations such as the BXDs [39] [40]. For example, AD estimates tend to have an *h_2_* close to 0.25, but, with eight replicates, the corresponding H2RIx̅ is 0.72, and there is a significant boost in power to detect effects (see power calculator power.genenetwork.org) To calculate empirical confidence intervals of heritabilities, we used two methods. First, *h_2_* values were recalculated, omitting one strain at a time. If heritability decreases when a particular strain is removed, the result indicates that the strain contributed a disproportionate fraction of V_a_. The second method involved omitting four random individuals from the analysis and recalculating heritabilities. This process was performed 10,000 times to provide an empirical estimate of the distribution of possible heritabilities using all four strains. Note that heritability estimates depend on the choice of strains and many potential environmental and technical confounds. When environmental variance is minimized, heritability will be higher. When technical variance is high, heritabilities will be depressed.

## DATA AND CODE AVAILABILITY

Scalar data for each specimen were aggregated into a single Excel sheet which contains two separate lists—left and right sides—for all 166 labels followed by volume, mean values of FA, AD, and RD for each sub region. Volume data for each sub region of a given strain were normalized to the total brain volume of that strain. The data package for each specimen consists of 51 unregistered 3D MRI volumes; a registered 4D volume; scalar images; mean values from the 166 regions for the scalars; tractography files from DSI studio; and connectomes. The aggregated data set is >500 GB. Representative data are available from our website (https://civmvoxport.vm.duke.edu/voxbase/studyhome.php?studyid=402). Access for additional data is provided by request at the same URL.

