## Supplemental Figures for "Heritability of the Mouse Brain Connectome"

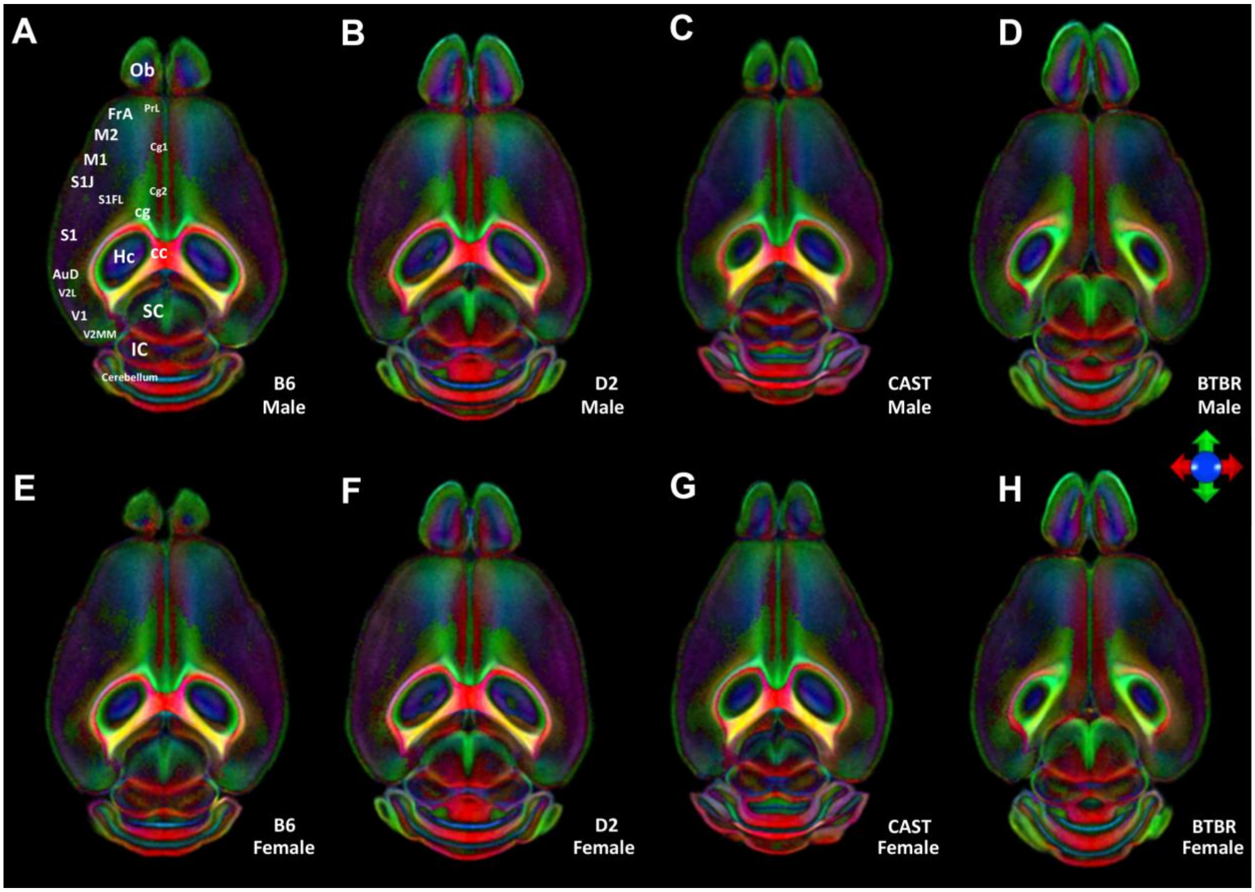


**Figure S1: Horizontal color FA images from the average (n=4) male and average female (n=4).** Images are color-coded to highlight the direction of fiber tracts (the principal eignenvector) in anterior commissure. **A)** B6, **B)** D2, **C)** CAST, **D)** BTBR. The callosum does not develop in BTBR. Color has been added by encoding the direction of the principal eigenvector from the diffusion tensor: red is left/right, green is anterior/posterior, and blue is dorsal/ventral. Ob: olfactory bulb; FrA: Frontal association Cortex; PrL: prelimbic; M2: primary motor; M2: secondary motor; Cg1: cingulate-area 1; Cg2: Cingulate-area 2; cg: cingulum; S1J; primary somatosensory – jaw; S1FL: primary somatosensory – forelimb; S1: primary somatosensory; AuD: secondary auditory; Hc: hippocampus; cc: corpus callosum; V2L: secondary visual – lateral area; V1: primary visual; V2MM: secondary visual – mediomedialarea; SC: superior colliculus; IC: inferior colliculus.

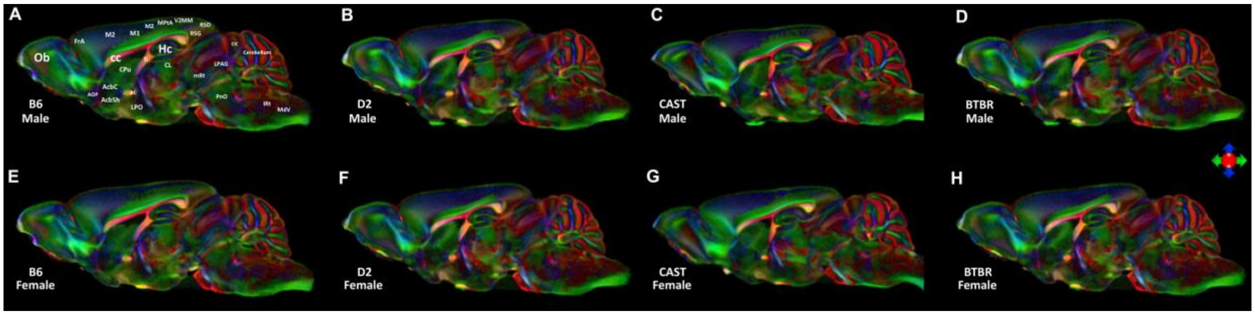


**Figure S2: Sagittal color FA images from the average (n=4) male and average female (n=4).** Images are color-coded to highlight the direction of fiber tracts (the principal eignenvector) in anterior commissure. **A)** B6, **B)** D2, **C)** CAST, **D)** BTBR. The callosum does not develop in BTBR. Color has been added by encoding the direction of the principal eigenvector from the diffusion tensor: red is left/right, green is anterior/posterior, and blue is dorsal/ventral. Ob: olfactory bulb; FrA: Frontal association Cortex; MO: Medial Orbital Cortex; S1J,S1ULP;: Primary Somatosensory Cortex;S2: secondary somatosensory cortex; AuD: secondary auditory cortex; TeA: Temporal association cortex; CEnt: caudomedial entorhinal cortex; HC: hippocampus; SC: superior colliculus; PAG: Periaquaductal grey; LGN: lateral geniculate nucleus; VT: ventral thalamic nuclei; LDNT: latero dorsal nucleus of thalamus; fi: Fimbria; cc: corpus callosum; SPT: Septum; Cpu: striatum;

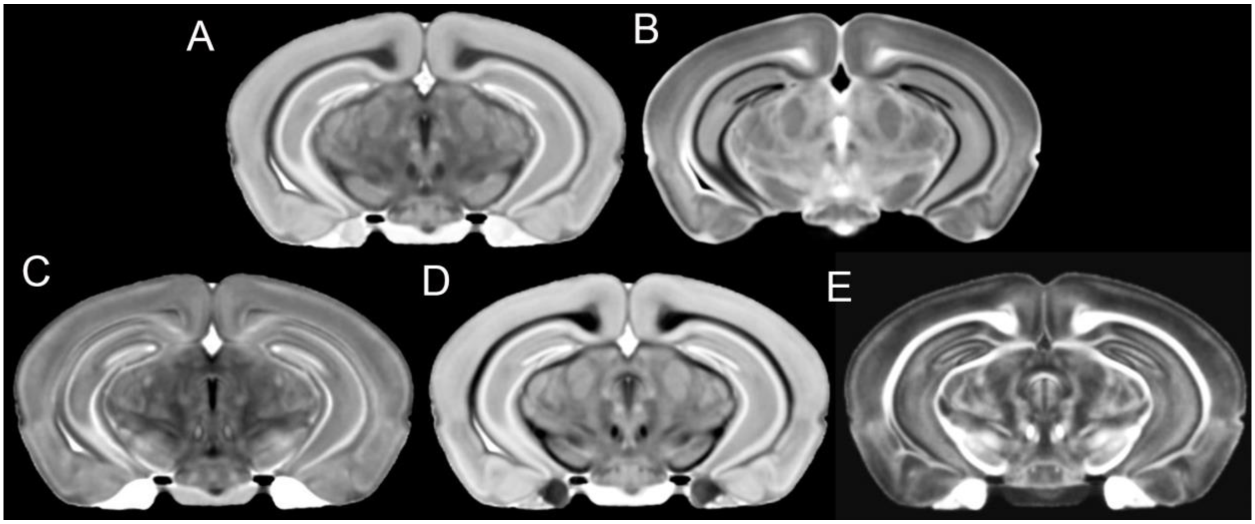


**Figure S3: Multiple scalar images are generated from the pipeline.** Shown here are A) ADC; B) DWI; C) AD; D) RD; E) FA. The complimentary contrast (e.g. DWI and FA) can be used to make label alignment more robust.

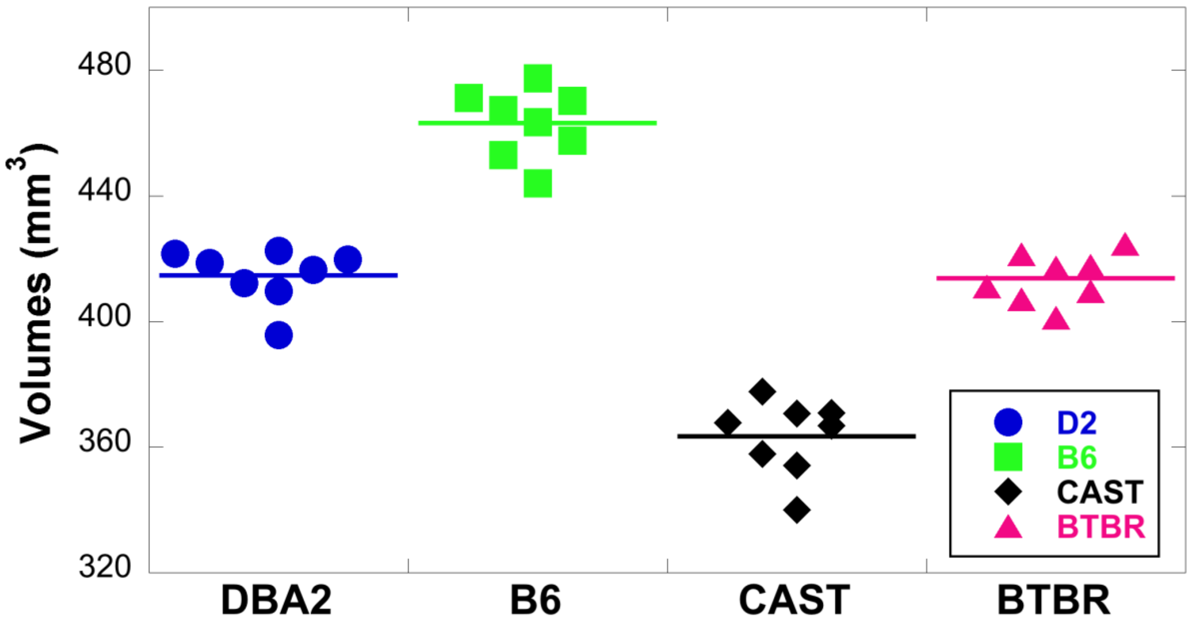


Figure S4: Total brain volume ($mm^3$) for eight animals each of four strains. Each point represents an individual, and the bar represents the mean for that strain.

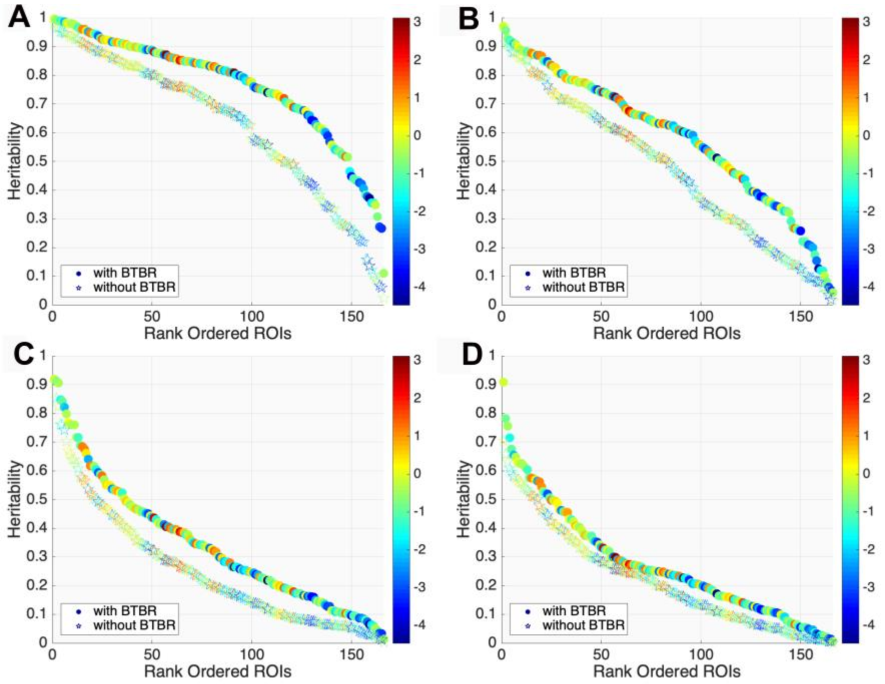


**Figure S5:** Broad sense heritability ($H_2$) vs rank ordered ROI for a) ROI mean volume; b) ROI mean fractional anisotropy; c) ROI mean radial diffusivity; d) ROI axial diffusivity. Circles show $H_2$ when all four strains are considered. Stars show $H_2$ when the BTBR is excluded. The color scale in each plot is a log for the value of that index (volume, fa,radial diffusivity, or axial diffusivity).

| Rank | Volume | | AD | | RD | | FA | |
|---|---|---|---|---|---|---|---|---|
| 1 | 5 | A29a__Cingulate_Cortex | 137 | n8__Vestibulocochlear_Nerve | 147 | cbw__Cerebellar_White_Matter | 147 | cbw__Cerebellar_White_Matter |
| 2 | 109 | CoN__Cochlear_Nucleus | 141 | sc__Spinocerebellar_Tract | 124 | cg__Cingulum | 27 | S1FL__Primary_Somatosensory_Cortex |
| 3 | 149 | A25__Cingulate_Cortex | 135 | sp5__Spinal_Trigeminal_Nerve | 118 | ac__Anterior_Commisure | 143 | vsc__Ventral_Spinocerebellar_Tract |
| 4 | 2 | A24aPrime__Cingulate_Cortex | 159 | PaR__Pararubral_Nucleus | 132 | bsc__Brachium_of_Superior_Colliculus | 19 | M1__Primary_Motor_Cortex |
| 5 | 143 | vsc__Ventral_Spinocerebellar_Tract | 126 | vhc__Ventral_Hippocampal_Commissure | 121 | cc__Corpus_Callosum | 132 | bsc__Brachium_of_Superior_Colliculus |
| 6 | 126 | vhc__Ventral_Hippocampal_Commissure | 96 | Vll__Ventral_Lateral_Lemniscus_Nucleus | 128 | fr__Fasciculus_Retroflexus | 146 | icp_oc_tz__Inferior_Cerebellar_Peduncle |
| 7 | 96 | Vll__Ventral_Lateral_Lemniscus_Nucleus | 120 | fi__Fimbria | 122 | fx__Fornix | 26 | S1DZ__Primary_Somatosensory_Cortex |
| 8 | 137 | n8__Vestibulocochlear_Nerve | 139 | lfp__Longitudinal_Fasciculus_of_Pons | 120 | fi__Fimbria | 28 | S1HL__Primary_Somatosensory_Cortex |
| 9 | 147 | cbw__Cerebellar_White_Matter | 13 | DLO__Dorsolateral_Orbital_Cortex | 146 | cp_oc_tz__Inferior_Cerebellar_Peduncle | 124 | cg__Cingulum |
| 10 | 63 | GP__Globus_Pallidus | 118 | ac__Anterior_Commisure | 84 | RPC__Red_Nucleus_Parvicellular | 153 | MiTg__Microcellular_Tegmental_Nucleus |
| 11 | 118 | ac__Anterior_Commisure | 60 | Preoptic_Telencephalon | 109 | CoN__Cochlear_Nucleus | 96 | Vll__Ventral_Lateral_Lemniscus_Nucleus |
| 12 | 92 | Den__Dentate_(Lateral)_Nucleus_of_Cerebellum | 146 | icp_oc_tz__Inferior_Cerebellar_Peduncle | 164 | Prerubral_Forel | 134 | ll__Lateral_Lemniscus |
| 13 | 123 | st__Stria_Terminalis | 17 | LO__Lateral_Orbital_Cortex | 131 | pc__Posterior_Commissure | 137 | n8__Vestibulocochlear_Nerve |
| 14 | 78 | RTN__Reticular_Nucleus_of_Thalamus | 109 | CoN__Cochlear_Nucleus | 108 | Tg__Tegmental_Nucleus | 80 | DpMe__Deep_Mesencephalic_Nuclei |
| 15 | 83 | SN__Substantia_Nigra | 150 | das__Dorsal_Acustic_Stria | 92 | Den__Dentate_(Lateral)_Nucleus_of_Cerebellum | 165 | PVG_of_Hypothalamus |
| 16 | 62 | SPT__Septum | 134 | ll__Lateral_Lemniscus | 95 | FasMed__Fastigial_Medial_Nucleus_of_Cerebellum | 41 | VO__Ventral_Orbital_Cortex |
| 17 | 1 | A24a__Cingulate_Cortex | 102 | m5__Motor_Root_of_Trigeminal_Nerve | 7 | A29c__Cingulate_Cortex | 13 | DLO__Dorsolateral_Orbital_Cortex |
| 18 | 105 | Raphe_Nucleus | 100 | CG__Central_Gray | 159 | PaR__Pararubral_Nucleus | 164 | Prerubral_Forel |
| 19 | 76 | LGN__Lateral_Geniculate_Nucleus | 131 | pc__Posterior_Commissure | 143 | vsc__Ventral_Spinocerebellar_Tract | 17 | LO__Lateral_Orbital_Cortex |
| 20 | 120 | fi__Fimbria | 41 | VO__Ventral_Orbital_Cortex | 106 | Pr5__Trigeminal_Sensory_Nucleus | 149 | A25__Cingulate_Cortex |

**Table S1: Rank order of the heritability of volume and DTI metrics.** A list of the top 20 heritable structures based on volume, AD, RD, and FA (ROI number and name)

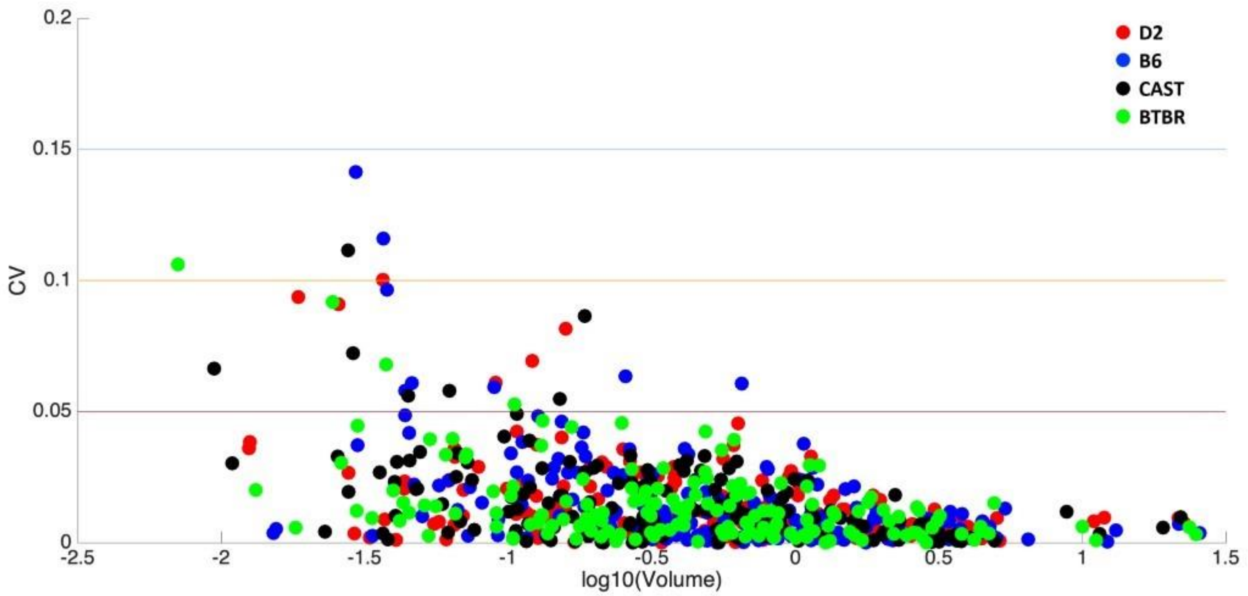


**Figure S6:** Coefficient of variation of left/right volume for D2, B6, CAST, and BTBR as a function of the mean ROI volumes.

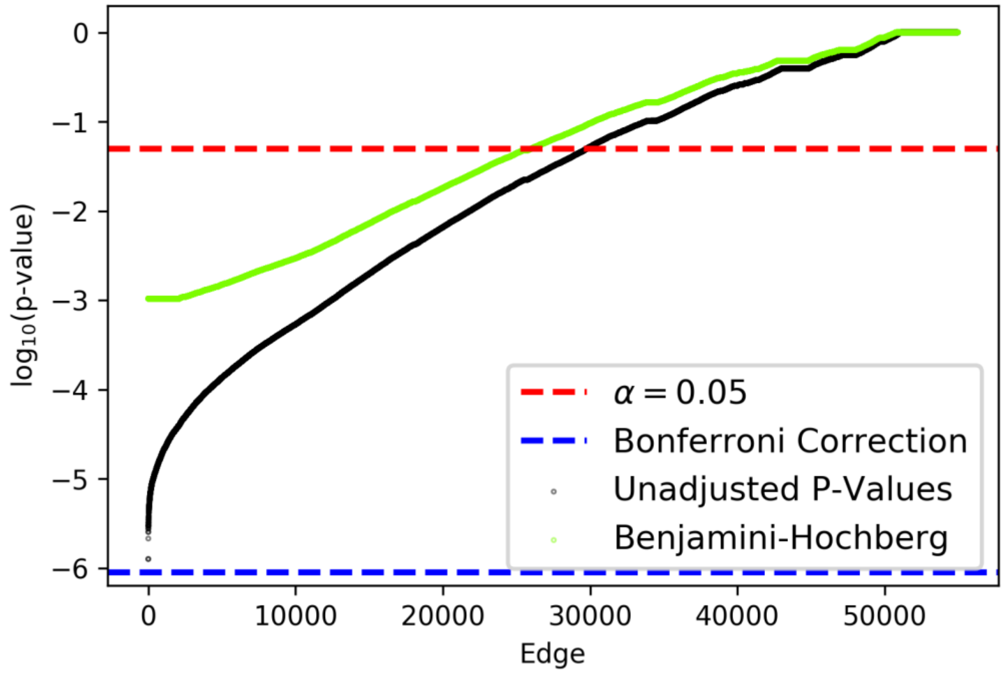


**Figure S7**: Nonparametric ANOVA p-values for the (n/2) edges. For each edge we have 4 samples of 8 univariate edge weights per sample – 4 genotypes, 8 animals per genotype. The x-axis is (n/2) edges, sorted by significance; the y-axis is log(p-value). Black dots are Kruskal-Wallis p-values; horizontal red line is p-value = 0.05; horizontal blueline is Bonferroni correction p-value 0.05/(n/2); green dots are Benjamini-Hochberg corrected p-values. We see that (1) most edges are significant at uncorrected p value = 0.05; (2) no edges are significant after Bonferroni correction; (3) most edges are significant with Benjamini-Hochberg correction.

| Rank | With BTBR | | Without BTBR | |
|---|---|---|---|---|
| 1 | 121 | cc__Corpus_Callosum | 127 | ic__Internal_Capsule |
| 2 | 249 | SN__Substantia_Nigra | 204 | V2L__Secondary_Visual_Cortex |
| 3 | 127 | ic__Internal_Capsule | 121 | cc__Corpus_Callosum |
| 4 | 286 | fi__Fimbria | 249 | SN__Substantia_Nigra |
| 5 | 299 | cp__Cerebral_Peduncle | 293 | ic__Internal_Capsule |
| 6 | 293 | ic__Internal_Capsule | 251 | MRN__Midbrain_Reticular_Nucleus |
| 7 | 66 | Acb__Accumbens | 252 | RI__Rostral_Linear_Nucleus |
| 8 | 287 | cc__Corpus_Callosum | 287 | cc__Corpus_Callosum |
| 9 | 62 | SPT__Septum | 77 | ZI__Zona_Incerta |
| 10 | 186 | M2__Secondary_Motor_Cortex | 83 | SN__Substantia_Nigra |
| 11 | 217 | Hc__Hippocampus | 133 | cp__Cerebral_Peduncle |
| 12 | 120 | fi__Fimbria | 66 | Acb__Accumbens |
| 13 | 178 | AuV__Secondary_Auditory_Cortex | 160 | Retro_Rubral_Field |
| 14 | 61 | Sthal__Subthalamic_Nucleus Sthal | 186 | M2__Secondary_Motor_Cortex |
| 15 | 65 | Amy__Amygdala | 132 | bsc__Brachium_of_Superior_Colliculus |
| 16 | 237 | VTA__Ventral_Tegmental_Area | 219 | Ect__Ectorhinal_Cortex |
| 17 | 133 | cp__Cerebral_Peduncle | 243 | ZI__Zona_Incerta |
| 18 | 292 | vhc__Ventral_Hippocampal_Commissure | 247 | SC__Superior_Colliculus |
| 19 | 289 | st__Stria_Terminalis | 202 | V1B__Primary_Visual_Cortex |
| 20 | 20 | M2__Secondary_Motor_Cortex | 229 | GP__Globus_Pallidus |

**Table S2: List of the nodes with the lowest p values from the manova comparisons.** The top 20 different nodes with BTBR and without BTBR. Left Hemisphere: ROI 1-166, Right Hemisphere: ROI 167-332.

| Value | Structure | Abbrev |
|---|---|---|
| 1 | A24a__Cingulate_Cortex,_Area_24a | A24a |
| 2 | A24aPrime__Cingulate_Cortex,_Area_24a_prime | A24aPrime |
| 3 | A24b__Cingulate_Cortex,_Area_24b | A24b |
| 4 | A24bPrime__Cingulate_Cortex,_Area_24b_prime | A24bPrime |
| 5 | A29a__Cingulate_Cortex,_Area_29a | A29a |
| 6 | A29b__Cingulate_Cortex,_Area_29b | A29b |
| 7 | A29c__Cingulate_Cortex,_Area_29c | A29c |
| 8 | A30__Cingulate_Cortex,_Area_30 | A30 |

| | | |
|---|---|---|
| 9 | A32__Cingulate_Cortex,_Area_32 | A32 |
| 10 | Au1__Primary_Auditory_Cortex | Au1 |
| 11 | AuD__Secondary_Auditory_Cortex,_Dorsal_Part | AuD |
| 12 | AuV__Secondary_Auditory_Cortex,_Ventral_Part | AuV |
| 13 | DLO__Dorsolateral_Orbital_Cortex | DLO |
| 14 | Fr3__Frontal_Cortex,_Area_3 | Fr3 |
| 15 | FrA__Frontal_Association_Cortex | FrA |
| 16 | Ins__Insular_Cortex | Ins |
| 17 | LO__Lateral_Orbital_Cortex | LO |
| 18 | LPtA__Lateral_Parietal_Association_Cortex | LPtA |
| 19 | M1__Primary_Motor_Cortex | M1 |
| 20 | M2__Secondary_Motor_Cortex | M2 |
| 21 | MO__Medial_Orbital_Cortex | MO |
| 22 | MPtA__Medial_Parietal_Association_Cortex | MPtA |
| 23 | PtPR__Parietal_Cortex,_Posterior_Area,_Rostral_Part | PtPR |
| 24 | S1__Primary_Somatosensory_Cortex | S1 |
| 25 | S1BF__Primary_Somatosensory_Cortex,_Barrel_Field | S1BF |
| 26 | S1DZ__Primary_Somatosensory_Cortex,_Dysgranular_Zone | S1DZ |
| 27 | S1FL__Primary_Somatosensory_Cortex,Forelimb_Region | S1FL |
| 28 | S1HL__Primary_Somatosensory_Cortex,_Hindlimb_Region | S1HL |
| 29 | S1J__Primary_Somatosensory_Cortex,_Jaw_Region | S1J |
| 30 | S1Sh__Primary_Somatosensory_Cortex,_Shoulder_Region | S1Sh |
| 31 | S1Tr__Primary_Somatosensory_Cortex,_Trunk_Region | S1Tr |
| 32 | S1ULp__Primary_Somatosensory_Cortex,_Upper_Lip_Region | S1ULp |
| 33 | S2__Secondary_Somatosensory_Cortex | S2 |
| 34 | TeA__Temporal_Association_Cortex | TeA |
| 35 | V1__Primary_Visual_Cortex | V1 |
| 36 | V1B__Primary_Visual_Cortex,_Binocular_Area | V1B |
| 37 | V1M__Primary_Visual_Cortex,_Monocular_Area | V1M |
| 38 | V2L__Secondary_Visual_Cortex,Lateral_Area | V2L |
| 39 | V2ML__Secondary_Visual_Cortex,_Mediolateral_Area | V2ML |
| 40 | V2MM__Secondary_Visual_Cortex,_Mediomedial_Area | V2MM |
| 41 | VO__Ventral_Orbital_Cortex | VO |
| 42 | CEnt__Caudomedial_Entorhinal_Cortex | CEnt |
| 43 | DIEnt__Dorsal_Intermediate_Entorhinal_Cortex | DIEnt |

| | | |
|---|---|---|
| 44 | DLEnt__Dorsolateral_Entorhinal_Cortex | DLEnt |
| 45 | MEnt__Medial_Entorhinal_Cortex | MEnt |
| 46 | VIEnt__Ventral_Intermediate_Entorhinal_Cortex | VIEnt |
| 47 | Cl__Claustrum | Cl |
| 48 | DCl__Dorsal_Claustrum | DCl |
| 49 | PLCo__Posterolateral_Cortical_Amygdaloid_Area | PLCo |
| 50 | VCl__Ventral_Claustrum | VCl |
| 51 | Hc__Hippocampus | Hc |
| 52 | DTT__Dorsal_Tenia_Tecta | DTT |
| 53 | Ect__Ectorhinal_Cortex | Ect |
| 54 | PaS__Parasubiculum | PaS |
| 55 | PRh__Perirhinal_Cortex | PRh |
| 56 | PrS__Presubiculum | PrS |
| 57 | Pir__Piriform_Cortex | Pir |
| 58 | APir__Amygdalopiriform_Transition_Area | APir |
| 59 | Hyp__Hypothalamus | Hyp |
| 60 | Preoptic_Telencephalon | Preoptic_Telencephalon |
| 61 | Sthal__Subthalamic_Nucleus | Sthal |
| 62 | SPT__Septum | SPT |
| 63 | GP__Globus_Pallidus | GP |
| 64 | Cpu__Striatum | Cpu |
| 65 | Amy__Amygdala | Amy |
| 66 | Acb__Accumbens | Acb |
| 67 | BNst__Bed_Nucleus_of_the_Stria_Terminalis | BNst |
| 68 | Vpal__Ventral_Pallidum | Vpal |
| 69 | PAG__Periaquaductal_Grey | PAG |
| 70 | APT__Anterior_Pretectal_Nucleus | APT |
| 71 | VTA__Ventral_Tegmental_Area | VTA |
| 72 | Thalamus_Rest | Thalamus_Rest |
| 73 | VT__Ventral_Thalamic_Nuclei | VT |
| 74 | LDNT__Latero_Dorsal_Nucleus_of_Thalamus | LDNT |
| 75 | MGN__Medial_Geniculate_Nucleus | MGN |
| 76 | LGN__Lateral_Geniculate_Nucleus | LGN |
| 77 | ZI__Zona_Incerta | ZI |
| 78 | RTN__Reticular_Nucleus_of_Thalamus | RTN |
| 79 | Sub_PP__Subbrachial_Nucleus_and_Peripeduncular_Nucleus | Sub_PP |
| 80 | DpMe__Deep_Mesencephalic_Nuclei | DpMe |
| 81 | SC__Superior_Colliculus | SC |

| | | |
|---|---|---|
| 82 | IC__Inferior_Colliculus | IC |
| 83 | SN__Substantia_Nigra | SN |
| 84 | RPC__Red_Nucleus_Parvicellular | RPC |
| 85 | MRN__Midbrain_Reticular_Nucleus | MRN |
| 86 | RI__Rostral_Linear_Nucleus | RI |
| 87 | Cn__Cuneiform_Nucleus | Cn |
| 88 | PrCn__Precuneiform_Nucleus | PrCn |
| 89 | Brain_Stem_Rest | Brain_Stem_Rest |
| 90 | Iped__Interpeduncular_Nucleus | Iped |
| 91 | 1Cb-10Cb__Cerebellar_Cortex | 1Cb-10Cb |
| 92 | Den__Dentate_(Lateral)_Nucleus_of_Cerebellum | Den |
| 93 | Int__Interposed_Nucleus_of_Cerebellum | Int |
| 94 | FasDL__Fastigial_Medial_Dorsolateral_Nucleus_of_Cerebellum | FasDL |
| 95 | FasMed__Fastigial_Medial_Nucleus_of_Cerebellum | FasMed |
| 96 | Vll__Ventral_Lateral_Lemniscus_Nucleus | Vll |
| 97 | ParB__Parabrachial_Nucleus | ParB |
| 98 | KF__Parabrachial_Medial_Nucleus_and_Koelliker_Fuse_Nucleus | KF |
| 99 | PCRt_Pr5__Parvicellular_Reticular_Nucleus_and_Principal_Sensory_Trigeminal_Nucleus | PCRt_Pr5 |
| 100 | CG__Central_Gray | CG |
| 101 | Su5__Pedunculotegmental,_Medial_Paralemniscial,_and_Supratrigemnial_Nuclei | Su5 |
| 102 | m5__Motor_Root_of_Trigeminal_Nerve | m5 |
| 103 | 5N__Trigeminal_Motor_Nucleus | 5N |
| 104 | PnC_PnO_PnV__Pontine_Reticular_Nucleus | PnC_PnO_PnV |
| 105 | Raphe_Nucleus | Raphe_Nucleus |
| 106 | Pr5__Trigeminal_Sensory_Nucleus | Pr5 |
| 107 | Dorsal_Tegmentum | Dorsal_Tegmentum |
| 108 | Tg__Tegmental_Nucleus | Tg |
| 109 | CoN__Cochlear_Nucleus | CoN |
| 110 | Pn__Pontine_Nucleus | Pn |
| 111 | RTg__Reticulotegmental_Nucleus_of_Pons | RTg |
| 112 | Olivary_Complex | Olivary_Complex |
| 113 | PnRt__Pontine_Reticular_Nucleus | PnRt |
| 114 | Sp5__Spinal_Trigeminal_Nucleus | Sp5 |
| 115 | Ve__Vestibular_Nuclei | Ve |
| 116 | Gi__Gigantocellular_Reticular_Nucleus | Gi |
| 117 | Cu__Cuneate_Nucleus | Cu |

| | | |
|---|---|---|
| 118 | ac__Anterior_Commisure | ac |
| 119 | ot__Optic_Tracts | ot |
| 120 | fi__Fimbria | fi |
| 121 | cc__Corpus_Callosum | cc |
| 122 | fx__Fornix | fx |
| 123 | st__Stria_Terminalis | st |
| 124 | cg__Cingulum | cg |
| 125 | lo__Lateral_Olfactory_Tract | lo |
| 126 | vhc__Ventral_Hippocampal_Commissure | vhc |
| 127 | ic__Internal_Capsule | ic |
| 128 | fr__Fasciculus_Retroflexus | fr |
| 129 | sm__Stria_Medularis | sm |
| 130 | mt__Mamillothalamic_Tract | mt |
| 131 | pc__Posterior_Commissure | pc |
| 132 | bsc__Brachium_of_Superior_Colliculus | bsc |
| 133 | cp__Cerebral_Peduncle | cp |
| 134 | ll__Lateral_Lemniscus | ll |
| 135 | sp5__Spinal_Trigeminal_Nerve | sp5 |
| 136 | py__Pyramidal_Tract | py |
| 137 | n8__Vestibulocochlear_Nerve | n8 |
| 138 | n7__Facial_Nerve | n7 |
| 139 | lfp__Longitudinal_Fasciculus_of_Pons | lfp |
| 140 | mlf_ts__Medial_Longitudinal_Fasciculus_and_Tectospinal_Tract | mlf_ts |
| 141 | sc__Spinocerebellar_Tract | sc |
| 142 | ml__Medial_Lemniscus | ml |
| 143 | vsc__Ventral_Spinocerebellar_Tract | vsc |
| 144 | mcp__Middle_Cerebellar_Peduncle | mcp |
| 145 | scp__Superior_Cerebellar_Peduncle | scp |
| 146 | icp_oc_tz__Inferior_Cerebellar_Peduncle | icp_oc_tz |
| 147 | cbw__Cerebellar_White_Matter | cbw |
| 148 | LV__Lateral_Ventricle | LV |
| 149 | A25__Cingulate_Cortex,_Area_25 | A25 |
| 150 | das__Dorsal_Acustic_Stria | das |
| 151 | Post__Postsubiculum | Post |
| 152 | 4V__Ventricular_System_4thVentricle | 4V |
| 153 | MiTg__Microcellular_Tegmental_Nucleus | MiTg |
| 154 | APT__Pretectal_Nucleus | Pretectal_Nucleus |

| | | |
|---|---|---|
| 155 | LDVL__Latero_Dorsal_Thalamic_Nucleus_Ventro_Lateral | LDVL |
| 156 | LP__Latero_Posterior_Nuclei_of_Thalamus | LP |
| 157 | Athal__Anterior_Thalamic_Nuclei | Athal |
| 158 | RMC__Red_Nucleus_Magnocellular | RMC |
| 159 | PaR__Pararubral_Nucleus | PaR |
| 160 | Retro_Rubral_Field | Retro_Rubral_Field |
| 161 | CSF | CSF |
| 162 | Intermediate_Reticular_Nucleus | Intermediate_Reticular_Nucleus |
| 163 | PHD_PaMP_Post_and_Lateral_Hypothalamus | PHD_PaMP_Post_and_Lateral_Hypothalamus |
| 164 | Prerubral_Forel | Prerubral_Forel |
| 165 | PVG_of_Hypothalamus | PVG_of_Hypothalamus |
| 166 | BLA_Basalateral_Amygdala | BLA_Basalateral_Amygdala |

**Table S3:** WHS labels, abbreviations and common names adapted from Calabrese et al [20].

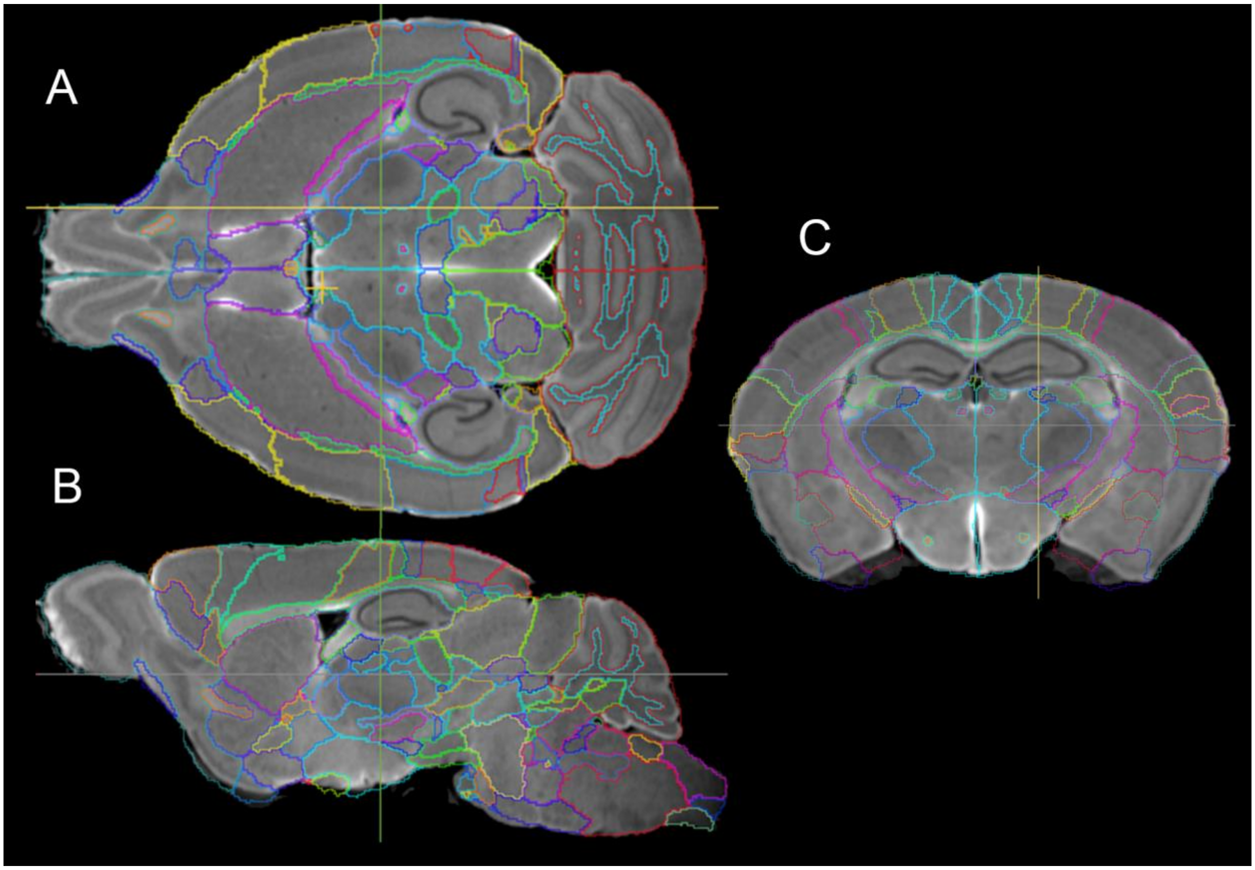


**Figure S8:** a) Dorsal; b) Sagital; c) Coronal sections of a C57 DWI with labels superimposed demonstrates their isotropic nature.

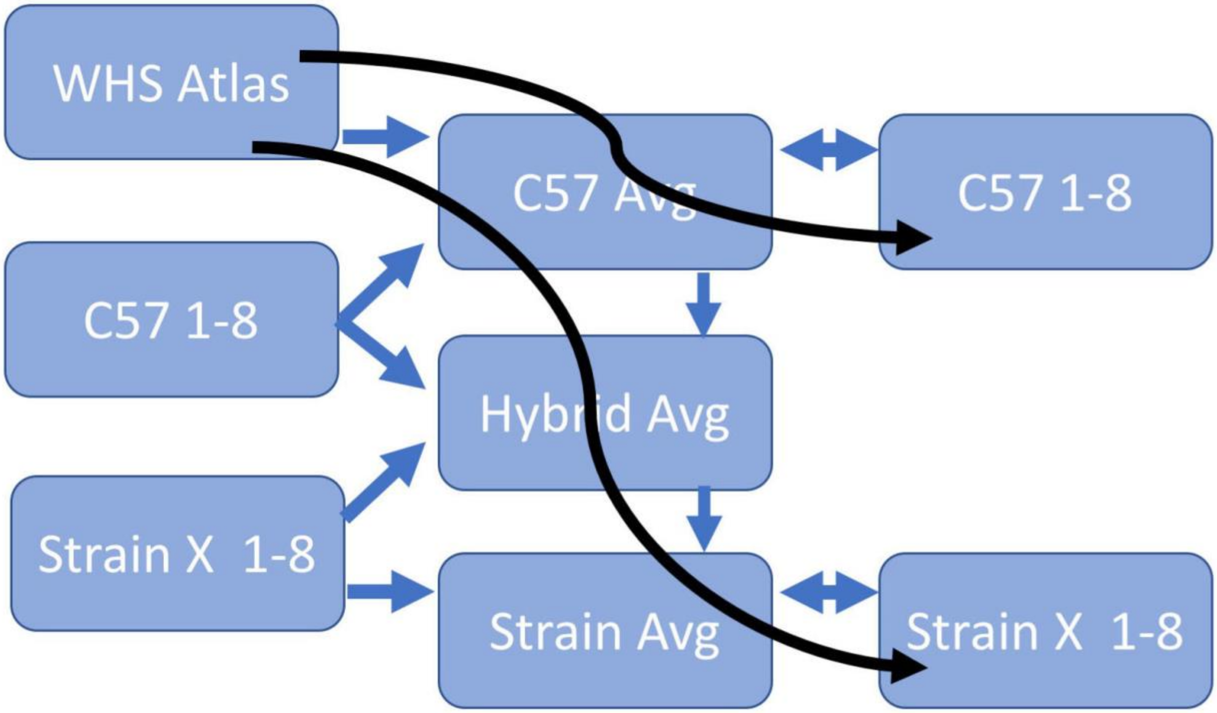


**Figure S9:** Mapping 3D labels from the WHS Atlas into the B6 mouse is straightforward. But mapping the same (B6) labels to the other strains requires an intermediate hybrid average. The individual image sets from each strain are then mapped into the strain average. The mapping is inverted leaving the WHS labels on each specimen in every atlas.

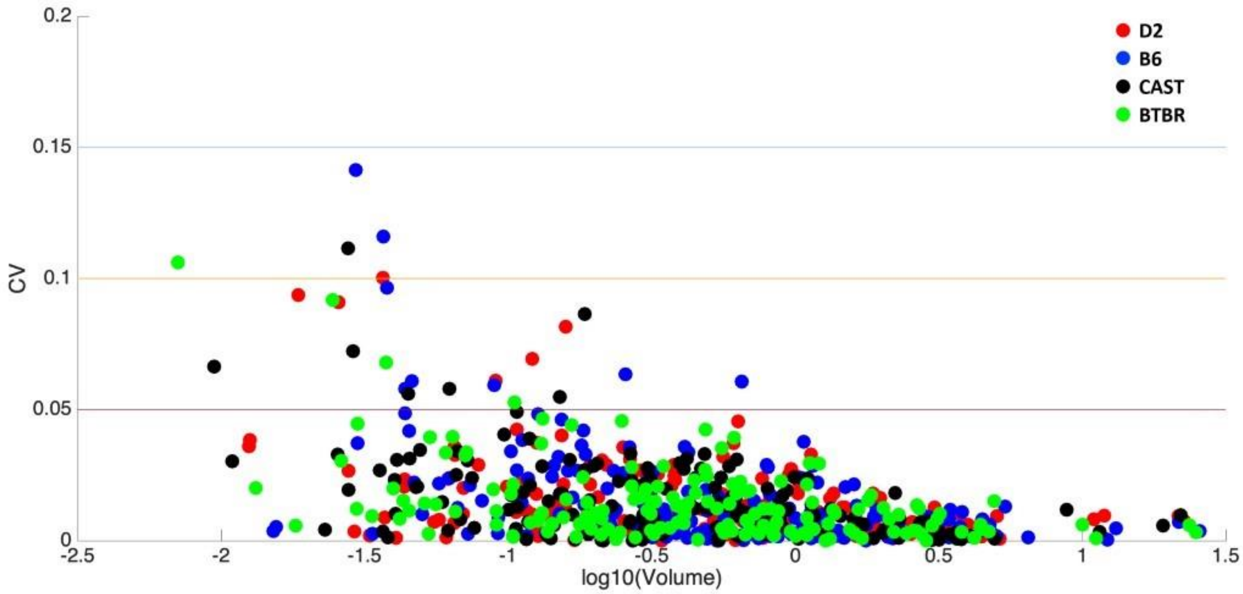

**Figure S10** Plot of the coefficient of variation as a function of substructure volume.
